# Developmental fidelity imposed by the RGL-1 “Balanced Switch” mediating opposing signals

**DOI:** 10.1101/381012

**Authors:** Hanna Shin, Christian Braendle, Kimberly B. Monahan, Rebecca E.W. Kaplan, Tanya P. Zand, F. Sefakor Mote, DeSean R. Craig, Eldon Peters, David J. Reiner

**Author notes:** These authors contributed equally to this study (Hanna Shin, Christian Braendle, Kimberly B. Monahan, Rebecca E.W. Kaplan). To whom correspondence should be addressed: David J. Reiner.

## Abstract

The six *C. elegans* vulval precursor cells (VPCs) are induced to form the 3^0^-3^0^-2^0^-Γ-2^0^-3° pattern of cell fates with high fidelity. In response to EGF signal, the LET-60/Ras-LIN-45/Raf-MEK-2/MEK-MPK-1/ERK canonical MAP kinase cascade is necessary to induce 1° fate and synthesis of DSL ligands. In turn, LIN-12/Notch signal is necessary to induce neighboring cells to become 2°. We previously showed that, in response to lower dose of EGF signal, the modulatory LET-60/Ras-RGL-1/RalGEF-RAL-1/Ral signal promotes 2° fate in support of LIN-12. In this study we identify two key differences between RGL-1 and RAL-1 functions. First, deletion of RGL-1 confers no overt developmental defects, while previous studies showed RAL-1 to be essential for viability and fertility. From this observation we hypothesize that the developmentally essential functions of RAL-1 are independent of upstream activation. Second, RGL-1 plays opposing and genetically separable roles in VPC fate patterning. RGL-1 promotes 2° fate via canonical GEF-dependent activation of RAL-1 and 1° fate via a non-canonical GEF-independent activity. Our genetic epistasis experiments are consistent with RGL-1 functioning in the modulatory 1°-promoting AGE-1/PI3-Kinase-PDK-1-AKT-1 cascade. Additionally, animals without RGL-1 experience 15-fold higher rates of VPC patterning errors compared to the wild type. Yet VPC patterning in RGL-1 deletion mutants is not more sensitive to environmental perturbations. We propose that RGL-1 functions as a “Balanced Switch” that orchestrates opposing 1°- and 2°-promoting modulatory cascades to decrease inappropriate fate decisions. We speculate that such switches are broadly conserved but mostly masked by paralog redundancy or essential genes.

## Introduction

Developmental patterning of the *C. elegans* vulva precursor cell (VPC) fates is a textbook system for analysis of cell-cell signaling. The vulva develops from six equipotent VPCs, P3.p through P8.p, which are induced to assume a 3°-3°-2°-1°-2°-3° pattern of cells fates. The anchor cell (AC) in the somatic gonad produces the LIN-3/EGF inductive signal (Fig. 1A). Historically, two models for VPC fate patterning have been advanced. The Morphogen Gradient Model posits that it is the position of each VPC in the LIN-3 gradient that dictates its fate. Notably, intact but isolated VPCs (the other five were killed by laser beam) could assume 1°, 2° or 3° fates, depending on their distance from the AC. Modulation of EGF or EGFR dose could control these outcomes (1–4). Thus, these were consistent with a graded inductive signal.

**Figure 1:**
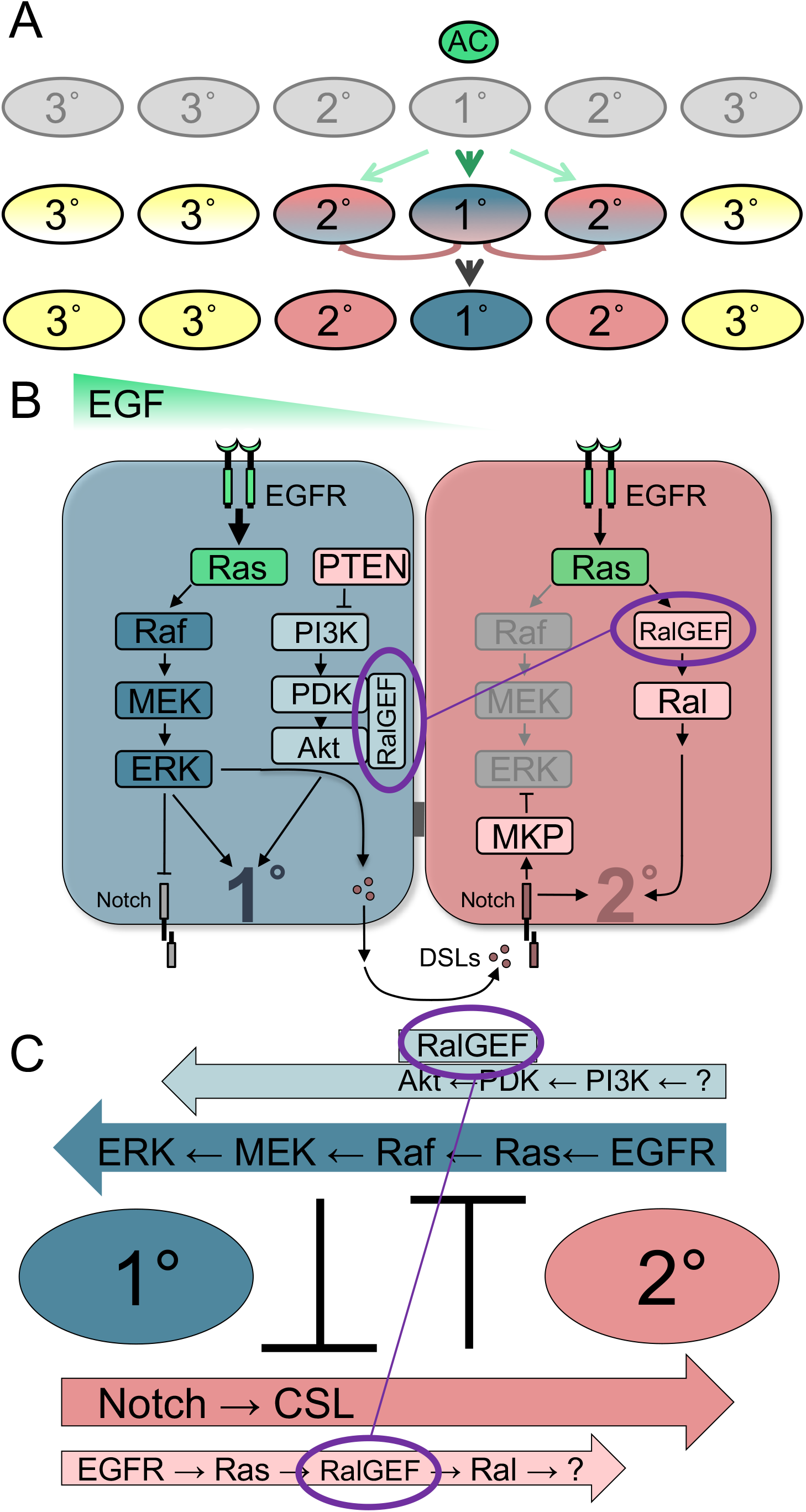
Schematics of VPC fate patterning and its signaling network. **A)** Initially equipotent VPCs are induced by the Anchor Cell (AC) to assume the 3°-3°-2°-1°-2°-3° pattern of fates (anterior-to-posterior), based on their position relative to the AC (see Introduction). Over time, induced VPCs progress from naïve to initially specified to terminally committed to their fates, represented by equipotent and uninduced (gray) progressing through initially specified (hybrid colors with one color dominant) to terminally committed to 1° (blue), 2° (rose) or 3° (yellow) fates. Yet the precise time course, molecular steps and network re-wiring events required for this progression are still unclear. **B)** The synthesis of the Sequential Induction, Morphogen Gradient, and Mutual Antagonism models of the VPC patterning signal transduction network, with the hypothesized RalGEF Balanced Switching mechanism superimposed. Names of human protein orthologs are used for ease of understanding between diverse experimental systems: EGF = LIN-3, EGFR = LET-23, Ras = LET-60, Raf = LIN-45, MEK = MEK-2, ERK = MPK-1, DSLs = DSL ligands, Notch = LIN-12, RalGEF = RGL-1, Ral = RAL-1, MKP (MAP kinase/ERK phosphatase) = LIP-1, PTEN = DAF-18, PI3K = AGE-1, PDK = PDK-1, Akt = AKT-1. Necessary and sufficient cascades are in dark colors (dark blue for 1°-promoting LET-23/EGFR-LET-60/Ras-LIN-45/Raf-MEK-2/MEK-MPK-1/ERK; dark rose for LIN-12/Notch-CSL transcriptional complex, CSL not pictured). Modulatory cascades are shown in lighter colors (light blue for 1°-promoting AGE-1/PI3K-PDK-1/PDK-AKT-1/Akt, with light rose for inhibitory DAF-18/PTEN lipid phosphatase; light rose for 2°-promoting RGL-1/RalGEF-RAL-1/Ral). Green represents proteins capable of promoting 1° or 2° fate, like LIN-3/EGF, LET-23/EGFR and LET-60/Ras, depending on signal dose and as yet unknown factors. Mutual antagonism operates by excluding potentially contradictory signals from initially specified VPCs: in presumptive 1 ° cells, LIN-12/Notch is internalized and degraded (gray), while in presumptive 2° cells MPK-1/ERK activation is repressed by transcriptional activation of LIP-1/MKP/DUSP/Erk phosphatase (and down-regulation of ral-1 promoter activity, not illustrated; (20)). The putative RGL-1/RALGEF Balanced Switch is circled in purple both in presumptive 1° and presumptive 2° cells, and connected by a thin purple line to represent a hypothetical switch is its signaling activity. **C)** A wiring diagram of the naive 1°/2° VPC patterning signaling network, illustrating parallel and anti-parallel signals, with essential signals in dark blue and rose, and modulatory signals in light blue and rose. Data support RGL-1/RalGEF functioning in antagonistic 1°-promoting (non-canonical) and 2°-promoting (canonical) cascades. Deletion of *rgl-1* would perturb both modulatory cascades but not alter the balance of 1°- and 2°-promoting signals, even in sensitized backgrounds.

In contrast, the Sequential Induction Model was posited upon identification and molecular genetic analysis of genes that are necessary and sufficient for the 1°- and 2°-promoting VPC fate patterning signals. LIN-3 signals via the LET-23/EGFR-LET-60/Ras-LIN-45/Raf-MEK-2/MEK-MPK-1/ERK canonical MAP kinase cascade to induce 1°fate (Fig. 1B; 5). LET-23 is necessary for 1° but not 2° fate (6, 7). However, these mosaic analyses did not address whether LET-23/EGFR might transduce a lower dose contribution to 2° fate induction, consistent with the Morphogen Gradient Model.

The Sequential Induction Model was further supported by the discovery of redundant DSL ligands, produced by presumptive 1° cells, that induce neighboring VPCs via LIN-12/Notch to become 2° (8, 9). LIN-12 is necessary and sufficient for 2° fate (10) and controls transcriptional client genes that were partially defined through computational means (Fig. 1B; 11-13).

Vulval induction is a stepwise progression: VPC fates are initially specified and later VPCs become committed (Fig. 1A; 14). Initial specification is accompanied by alterations of expression of certain signaling genes. Generally, expression of these is either uniformly absent or uniformly present in naive VPCs. Yet by the first cell division, several genes initiate lineage-specific expression patterns. Some of these transcriptional changes contribute to the Mutual Antagonism model, whereby contradictory signals are excluded from cells committing to cell fate. For example, the LIP-1/ERK phosphatase, a LIN-12 transcriptional client gene, is expressed in initially specified 2°s to restrict inappropriate ERK activation (11, 12). Conversely, LIN-12, via mechanisms not entirely understood, is targeted for internalization and degradation in initially specified 1° cells to restrict inappropriate 2°-promoting signal there (15–17).

Taken together, these many observations support the Sequential Induction model, yet fail to account for the experiments that implicate the action of an EGF gradient. The contradiction between these two models remained unresolved for 16 years. (18). We reconciled these potentially contradictory models by identifying the mechanism by which the 2°-promoting activity of the EGF gradient was transduced. We concluded that both models are true: initial sequential induction establishes the specification of the VPC pattern, and that overlaid on this pattern is a spatial EGF gradient that reinforces the initial pattern. To interpret different doses of EGF signal, LET-60/Ras dynamically switches effectors during VPC fate patterning, from LIN-45/Raf signaling through the canonical MEK-2/MEK-MPK-1ERK MAP kinase cascade to promote 1° fate, to RGL-1/RalGEF-RAL-1/Ral signaling to promote 2° fate (19, 20).

We have further found that RAL-1 promotes 2° fate through EXOC-8/Exo84, a subunit of the heterooctomeric exocyst complex, a known downstream oncogenic Ral signaling intermediary in mammalian cells. The GCK-2 MAP4 kinase and a downstream PMK-1/p38 MAP kinase cascade is necessary and sufficient for Ral-dependent increased induction of 2° cells, in support of the necessary and sufficient LIN-12 (Shin *et al.,* in press).

More recently, this VPC fate patterning system has been exploited to investigate more sophisticated questions in developmental biology. Vulval development has been used to compare mechanisms of induction across nematode species both closely and distantly related to *C. elegans* (21–24). Additionally, VPC fate patterning has been used to assess heterogeneity in polymorphic wild *C. elegans* isolates (22, 25), and also developmental robustness in the face of environmental insult: the VPCs are pattered with 99.8% accuracy (26). But this accuracy is compromised by weak mutations in the signaling network that alter the delicate balance of 1 ° and 2° inductive signals (27–30). One potential extrapolation from these studies is that there are heretofore unidentified properties of signaling networks that increase fidelity and/or robustness.

Ras is the most mutated mammalian oncoprotein: more than a quarter of all tumors - - up to 95% of Pancreatic Ductal Adenocarcinomas (PDAC) – harbor activating mutations in a Ras gene (31). Three main oncogenic Ras effector cascades have been identified (Fig. S1). Two of these, the Raf-MEK-ERK canonical MAP kinase cascade and the PI3-Kinase-PDK-Akt cascade, are among the best-studied and most pharmacologically targeted cascades in all of biology (32–34). In contrast, the RalGEF-Ral effector signal is poorly characterized, though some argue it is at least as important for oncogenesis as Raf and PI3K cascades (35–40). Ras binds and activates RalGEF, an exchange factor that stimulates GTP loading on the Ral small GTPase. Ral is a Ras-like small GTPase subfamily, in the Ras family of the Ras superfamily, and is conserved throughout Metazoa (41). All Ras family members, including Ral, are similar structurally: they cycle between inactive GDP- and active GTP-bounds states, with the latter promoting effector binding. For most Ras family members, including mammalian RalA and RalB, this GDP/GTP cycle is controlled by activating GEFs and inactivating GAPs. And, despite their structural and primary sequence similarities, closely related GTPases in the same family diverge at the core effector binding sequence, in the Switch I region, that interfaces directly with effectors. Thus, Ral interacts with a very different set of effectors than does Ras (42).

Notably, the core effector-binding sequences of *C. elegans* LET-60 and RAL-1 are 100% identical to those of their *Drosophila* and mammalian counterparts (20, 41). Known orthologs of these effectors are also conserved across Metazoa, so the field has operated on the reasonable assumption that the different suites of effectors employed by Ras vs. Ral are conserved across species.

Mammals encode three Ras proteins (K-, N-, H-Ras), four GEFs with RA domains (RalGDS, RGL, RGL1, RGL2), two Rals (A, B), three Rafs (A-, B-, C-Raf) and three PI3Ks (a, β, δ). *C. elegans* and *Drosophila* each harbor single Ral- and RalGEF-encoding genes, as well as single Ras-, Raf-, and PI3K-encoding genes (Fig. S1B). As is frequently the case with the multiplicity of mammalian genes compared to single invertebrate genes, these multiple isoforms likely impose an additional layer of complexity in signaling networks and cell biological outputs (43).

All three oncogenic Ras effectors have been implicated in *C. elegans* VPC fate patterning. Thus, we know of essential and modulatory cascades promoting both 1° and 2° fate: the essential LET-60-LIN-45 cascade promotes 1° fate with support of the modulatory AGE-1-PDK-1-AKT-1 cascade, and the essential LIN-12/Notch cascade promoters 2° fate with support of the modulatory LET-60-RGL-1-RAL-1 cascade.

Our analysis pursues the unexpected finding that RGL-1/RalGEF is functionally is non-equivalent to RAL-1 in two distinct ways. First, while RAL-1 is essential for viability and fertility (20, 44), RGL-1 is not essential. Second, RGL-1 and RAL-1 are nonequivalent in VPC fate patterning. While RAL-1 functions as a simple intermediary to propagate the 2°-promoting LET-60-RGL-1-RAL-1 signal, RGL-1 additionally performs an opposing function, a putative 1°-promoting function, that offsets its canonical 2°-promoting function. As a consequence, deletion of RGL-1 has no net effect on the delicate balance of 1°- and 2°-promoting signals in our most sensitized mutant backgrounds. Using GEF-specific mutations and genetic bypass experiments, we show that the opposing functions of RGL-1 in VPC fate patterning are genetically separable and function cell autonomously in VPCs. In the context of mammalian studies that argue that RalGEF physically interacts with PDK and Akt and functions as a scaffold (45, 46), our genetic epistasis results are consistent with RGL-1 functioning as a scaffold for PDK-1 and AKT-1 in the modulatory 1°-promoting AGE-1/PI3K cascade. Our analysis raises the question of how activity in two apparently opposing cascades contribute to VPC fate patterning. We found no difference in response of two RGL-1 deletion mutants and the wild type to environmental insults. Yet error rate in VPC patterning was 15-fold higher in the *rgl-1* deletion mutants than in the wild type. We hypothesize that the two opposing activities of RGL-1, which tie together the two opposing 1° and 2°-promoting modulatory cascades in VPC fate patterning, are orchestrated to reduce the level of noise in the signaling network, and hence reduce the rate of ambiguous fates or mis-patterning events. We conjecture that these properties of RGL-1 identify a novel switch to coordinate modulatory signaling activities in the reinforcement stage of VPC fate patterning that leads to fate commitment, and hence increases fidelity of the developmental process.

## Results

### The *C. elegans* RalGEF ortholog, RGL-1, is non-essential

We previously described that *ral-1(tm2760)* mutant animals are sterile but otherwise wild type. Efforts to feed or inject dsRNA in the RNAi hypersensitive *rrf-3* background failed to phenocopy this sterility. Consistent with our depletion by bacterially mediated RNAi and injected dsRNA, *tm2760* abrogated the 2°-promoting activity of RAL-1 (20).

Subsequent analysis of *ral-1,* including the more recently isolated allele, *ral-1(tm5205),* led to the conclusion that RAL-1 function is maternally rescued and necessary for various facets of embryonic, post-embryonic, and germline development, including function of the PAR complex in cell polarity (44). This function is ascribed to a central role of RAL-1 in the exocyst complex, as described for mammals (47–52). Consistent with published results, we also observed that *ral-1(tm5205)* mutants become sickly in the fourth larval stage and become sterile adults, but that the VPCs are patterned normally (N = 52).

Mammalian RalA and RalB associate with Sec5 and Exo84 subunits of the heterooctomeric exocyst complex (47–52). This functional association with the exocyst complex is consistent with results in *C. elegans* (20, 44), through the C-terminal membrane-targeting of Ral localizing the exocyst to specific subcellular domains to support certain exocyst functions. Yet mammalian studies suggest that Ral associates with the exocyst in an activity-dependent manner (47–52). This model predicts that abrogation of GTP-loading, either through blockade of total RalGEF activity or Ral GTP-loading activity, should phenocopy loss of Ral and result in defective exocyst function. *C. elegans,* which encodes only a single RalGEF ortholog, orthologous to mammalian RA-domain containing proteins (20), provides a system to test this relationship between RalGEF and Ral *in vivo.*

Also described previously, RNAi depletion of *rgl-1* revealed a role in promoting 2° fate (20). We reproduced RNAi depletion of *rgl-1* with bacterially mediated and injected dsRNA in wild-type and *rrf-3* mutant backgrounds and observed no overall development and fertility. Thus, we were unable to resolve this question by depletion of gene product via RNAi, and proceeded to analyze *rgl-1* mutations not used in the previous study. Strikingly, four different deletion mutants of RGL-1 are superficially wild type (Table 1). By visual inspection, mutants lacking functional RGL-1 develop normally, are fertile, and can be grown indefinitely as homozygous mutant strains. One of these mutations, *fax-1(gm27),* deletes several neighboring genes on the X chromosome, including *rgl-1* (53). Using robust primer sets (see Methods; Figs. S6, S7), we failed to amplify *rgl-1* sequences from the *gm27* mutant animal.

**Table 1:** *rgl-1* and *ral-1* alleles and mutant phenotypes

| Allele | Lesion | Superficial Phenotype | Net VPC alteration<sup>a</sup> |
| --- | --- | --- | --- |
| *rgl-1(ok1921)* | In-frame deletion, GEF domain | Wild type | none |
| *rgl-1(tm2255)* | Out-of-frame deletion, GEF domain | Wild type | none |
| *fax-1(gm27)<sup>b</sup>* | *rgl-1* and 8 other genes deleted | Unc<sup>c</sup> | none |
| *rgl-1(gk275305)* | nonsense | Wild type | none |
| *rgl-1(gk275304)* | Missense, R361Q, GEF domain | Wild type | Decreased 2° |
| *ral-1(tm2760)* | Intron 3 deletion | Sterile<sup>d</sup> | Decreased 2°<sup>d</sup> |
| *ral-1(tm5205)* | Exons 2 and 3 deletion | Variable, Sterile<sup>e</sup> | Decreased 2°<sup>e</sup> |
| *ral-1(gk628801)* | Missense, R139H | Wild type | Decreased 2° |
| *ral-1(re179)* | mNeonGreen::3xFlag::rgl-1 | Wild type | none<sup>f</sup> |

<sup>a</sup> Alteration in network signaling inferred from phenotype in *let-60(n1046gf)* background
<sup>b</sup> Documented in (53).
<sup>c</sup> Deletion of *fax-1* causes an Uncoordinated phenotype (53).
<sup>d</sup> Intron 2 deletion breaks in the middle of the splice donor site, conferring sterility and reduced 2° induction (20).
<sup>e</sup> Exons 2 and 3 deletion removes GTPase domain sequences, confers sterility, delayed growth and reduced 2° induction (44), this study.
<sup>f</sup> No effect in *let-60(n1046gf)* background

We also previously published that nonsense mutants for the alpha or beta subunits of the heterodimeric RalGAP, HGAP-1 and HGAP-2, respectively, are also viable and fecund (54). The G26V putative activating mutation in RAL-1 also fails to confer developmental defects in an otherwise wild-type background (Shin *et al.,* in press). The observation that deletion of RGL-1/RalGEF or HGAP-1/2/RalGAP does not result in the same phenotype as deletion of RAL-1 raises the interesting possibility that Ral and RalGEF are functionally non-equivalent, contrary to the model derived from mammalian studies. Thus, our observations suggest that the role of RAL-1 in the exocyst is independent of GDP/GTP-bound state.

### Sensitized genetic backgrounds reveal nuances in signaling

Four signaling cascades – two central and two modulatory – control 1 ° and 2° fate induction. We present a schematic of the signaling cascades discussed in this study (Fig. 1B,C; Fig. S1). Core 1°- and 2°-promoting signals are detectable by direct mutation: since they are necessary and sufficient to induce their respective fates, mutational perturbation of them causes loss or gain of vulval cell types. In contrast, the role of the two modulatory cascades is not revealed through single mutant analysis, but rather requires sensitized genetic backgrounds. We propose that RGL-1/RalGEF participates in both modulatory cascades (see below).

To detect such signals, we use *let-60(n1046gf),* a moderately activating G13E mutation analogous to mutations found in some mammalian cancers (55). In this background, gain and loss of the RAL-1 2°-promoting signal resulted in decrease and increase of ectopic 1° cells, respectively. We have similarly used the *let-23*(*sa62*gf) activating mutation in the LET-23/EGFR, and for under-induced backgrounds used the reduced function *lin-45(n2506)* background (20). Through combined use of genetic principles of parallelism and epistasis, we are able to dissect these modulatory signals. We present these principles as a network circuitry diagram (Fig. 1C). We have exploited such techniques to delineate a 2°-promoting signaling cascade downstream of RAL-1 (Shin *et al.,* in press), and here use these techniques to similarly dissect the two roles of RGL-1/RalGEF in 2°- and 1°-promoting activities.

### RGL-1 performs a function in VPC cell fate patterning that opposes its canonical 2°-promoting function

We previously showed that depletion of *rgl-1* by RNAi revealed a role of RGL-1 in promoting 2° fate consistent with the established Ras>RalGEF>Ral signal in mammals (20). We reproduce these RNAi-based experiments here (Fig. 2A-C, E). Drawing from analysis of *ral-1(tm5205)* allele, which deletes RAL-1 sequences universally conserved in the small GTPase superfamily (44; our unpublished results), we analyzed the impact of *tm5205* on VPC fate patterning. We found that *ral-1(tm5205)* confers enhancement of 1° induction in the *let-60(n1046gf)* background (Fig. 2F).

**Figure 2.**
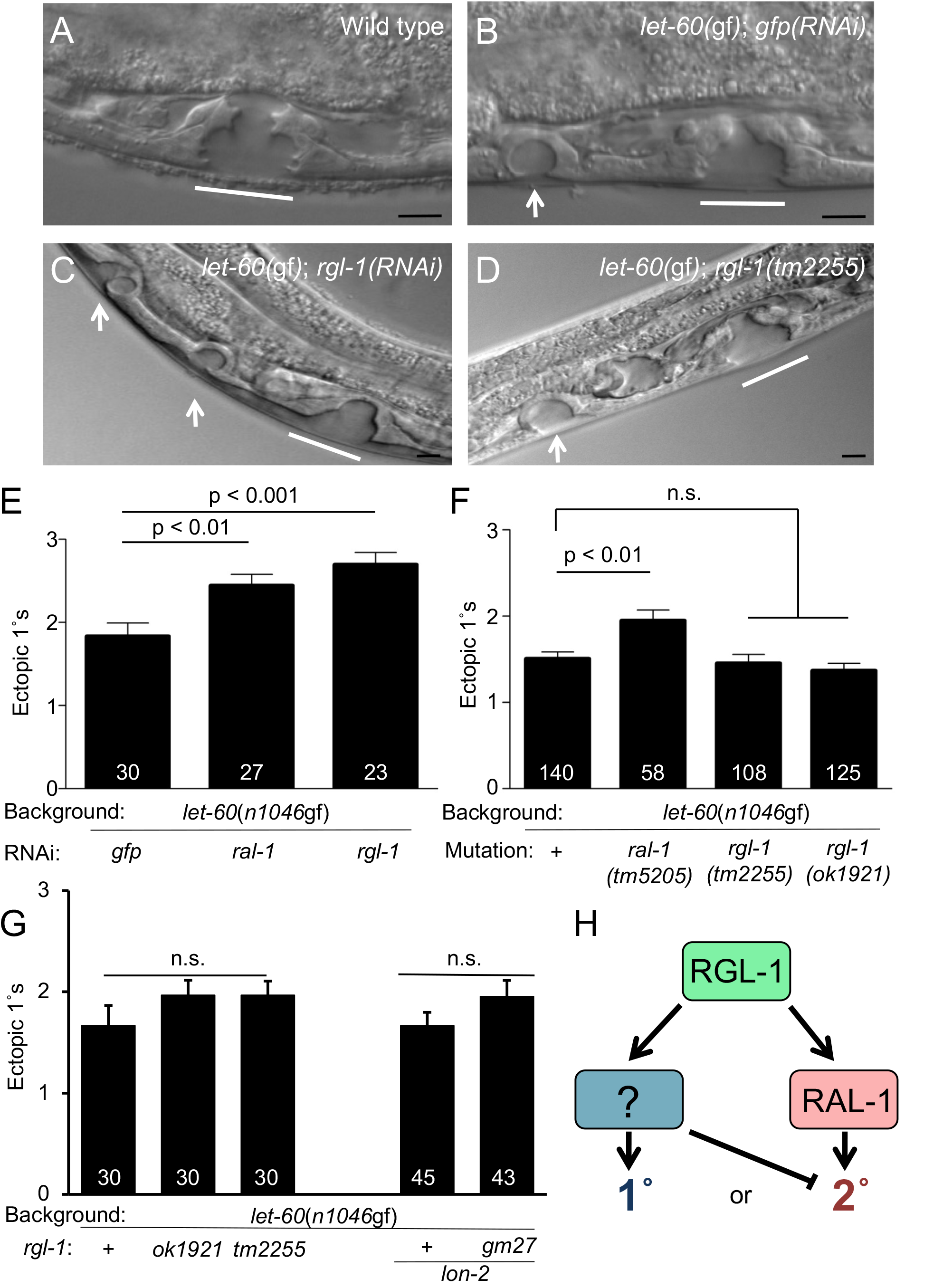
RGL-1 and RAL-1 are functionally non-equivalent in VPC patterning. 1000x photomicrographs of late L4 **A)** wild-type, **B)** *let-60(n1046gf); gfp(RNAi)* vs. 600 x photomicrographs of **C)** *let-60(n1046gf); rgl-1(RNAi),* and **D)** *let-60(n1046gf); rgl-1(tm2255)* animals. White lines = normal 2°-1°-2° L4 vulvae, white arrows = L4 ectopic 1° pseudovulvae. Black scale bars = 10 μm. Anterior is left and ventral down. **E)** RNAi depletion of *rgl-1* and *ral-1* enhance 1° induction relative to *gfp(RNAi)* control. Data are the mean ± standard error of the mean (SEM). For statistical reasons single, non-pooled assays are shown, and white numbers represent animals scored therein. Significance was calculated by Kruskal-Wallis, Dunn test. Data shown were scored concurrently and are representative of 4 independent assays (this study) and 6 prior independent assays (20). (The *let-60(n1046gf)* 1° induction baseline is consistently higher when grown on HT115 bacterially-mediated RNAi food compared to the standard 0P50; (20); Shin et al. in press.) **F)** Deletion of *ral-1* but not *rgl-1* enhances ectopic 1° induction by *let-* 60(n1046gf). Data shown are representative of 4 assays, each scored concurrently. **G)** Re-constructed strains show the same result: *ok1921, tm2255* and *gm27* deletions fail to significantly enhance ectopic 1° induction by *let-60(n1046gf).* Left: Three *n1046-* containing isolates, with and without *rgl-1* mutations and scored concurrently. The concurrently scored MT2124 1° induction baseline was not significantly different from outcrossed lines DV2214 and DV2215 (see Table S1), N = 30 for each, and from assays that showed the most deviation of double mutants from the single mutant, but are still not significantly different. Right: DV2251 *let-60(n1046gf); lon-2(e678)* vs. DV2252 *let-* 60(n1046gf); *lon-2(e678) gm27* animals scored concurrently but separate from the left group, representative of two assays. **H)** A general model for opposing RGL-1 GEF/RAL-1-dependent and -independent functions (green = bifunctional, blue = 1°-promoting, rose = 2°-promoting, see Fig. 1).

To our surprise, analysis of various strong loss or putative null *rgl-1* mutations in the *let-60(n1046gf)* background caused no net effect compared to the *let-60(n1046gf)* single mutant (Fig. 2D, F, G; see Table 1 and Fig 4A for *rgl-1* alleles), contrasting with depletion of *rgl-1* by injected or bacterially mediated RNAi. We replicated these results using a 1° fate reporter that indicates neighboring 1° cells in the *let-60(gf)* background (12, 20). Consistent with our previous results, *rgl-1(RNAi)* significantly increases the occurrence of neighboring 1° cells, but *rgl-1(tm2255)* does not (Fig S2A-E). Conscious that the *let-60(n1046gf)* mutant phenotype can drift and become more severe with protracted culturing, we used a stringent protocol for its analysis (20; Shin *et al.,* in press; see Methods). To test that these results are not specific to the *let-60(n1046gf)* sensitized background or a 1° over-inducing background, we also assessed the role of deleted *rgl-1* in under-induced background *lin-45(n2506)* and over-induced background *let-23(sa62gf)* (2, 20). *rgl-1(tm2255)* did not alter the balance between 1° and 2° induction (Fig. S2F,G), consistent with our results in the *let-60(n1046gf)* background.

Our results are consistent with the working model that RGL-1 performs an additional, Ral-independent function that antagonizes its canonical function, perhaps by promoting 1° fate. Consequently, deleting both functions results in no net change in the delicate balance between 1°-promoting and 2°-promoting signals in our sensitized genetic backgrounds. The discrepancy between RNAi- and mutational-based analyses is consistent with RGL-1 having different functional thresholds in level of gene product for the two opposing activities, which we have been unable to further investigate.

### The *rgl-1* transcriptional fusion is expressed in both 1° and 2° lineages

We previously described a transgenic *ral-1* transcriptional fusion that expressed GFP dynamically over the time course of VPC fate induction (20). To summarize published results, early in the 3^rd^ larval stage (L3), GFP was expressed consistently in all six VPCs. Later in L3, after induction, GFP expression was excluded from presumptive 1 ° cells while persisting in presumptive 2° cells. These observations provided a potentially critical mechanistic insight: by reducing inferred RAL-1 expression in presumptive 1° cells while retaining expression in presumptive 2° cells, the signaling network could attenuate inappropriate RAL-1 activation in presumptive 1° cells, thereby preventing potentially contradictory LET-60 signaling through RGL-1 -RAL-1 in presumptive 1° cells.

We similarly tested the expression pattern of a transgene harboring the *rgl-1* promoter transcriptional fusion to GFP, *sEx14985* (56). As with the *ral-1* promoter transcriptional fusion, the transgenic *rgl-1* reporter was expressed in all VPCs early in L3 (Fig. 3). But, unlike the *ral-1* transcriptional fusion, the *rgl-1* transcriptional fusion continued to express GFP in presumptive 1° cells throughout vulval patterning, proliferation and morphogenesis. Persistent GFP expression from the *rgl-1* promoter is consistent with both canonical Ral-dependent and non-canonical Ral-independent functions of RGL-1, were the opposing Ral-independent function 1°-promoting rather than 2°-promoting.

**Figure 3:**
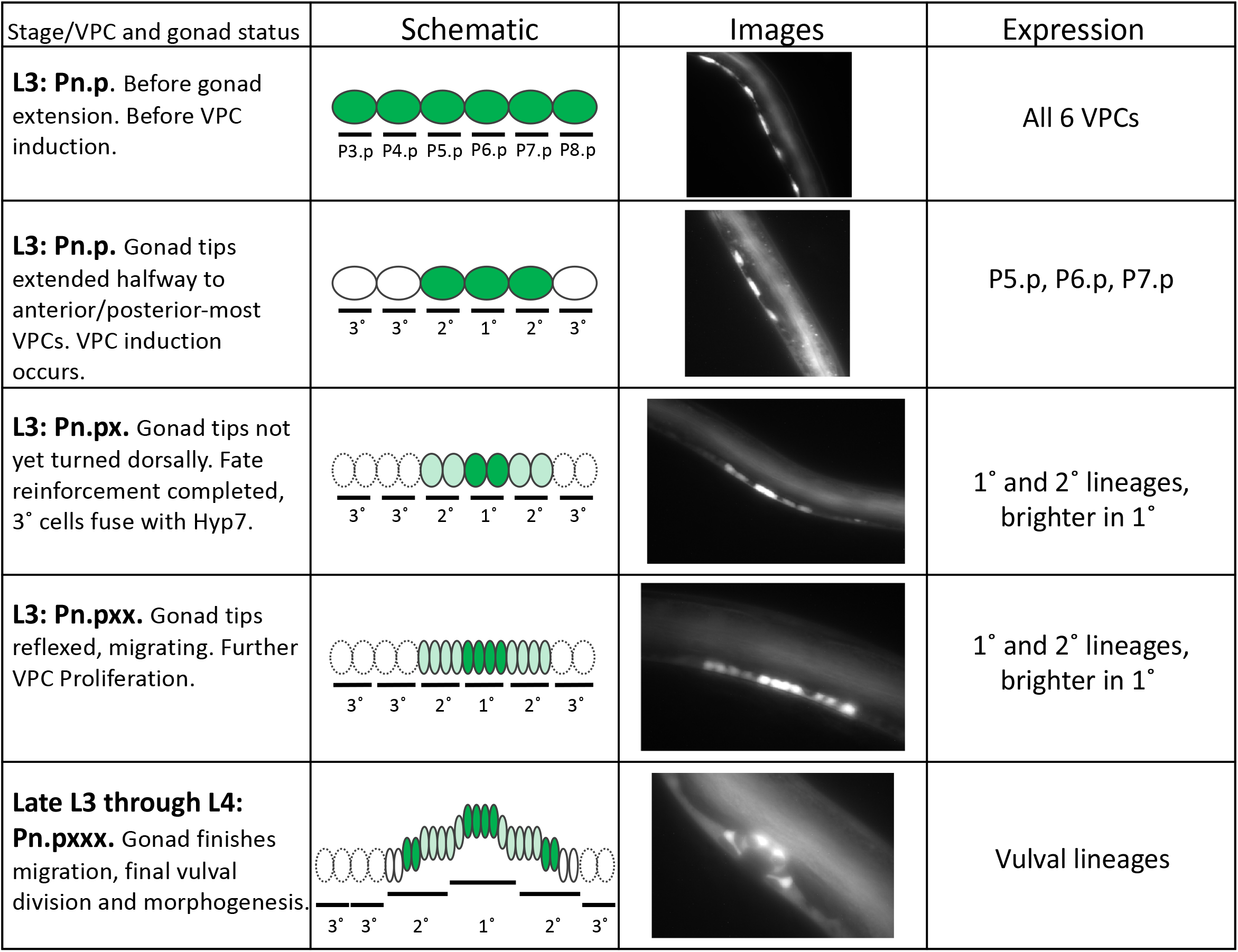
VPC expression pattern of the *rgl-1* transcriptional GFP fusion over time. We used a combination of DIC analysis of VPCs and migration of the gonadal distal cells for staging to characterize the dynamic pattern of GFP expression from *sEx14985* (56) over time. Initial expression in naïve VPCs is uniform. Around the time of induction, expression is restricted to presumptive vulval lineages. Later expression, after the first cell division, remains higher in 1° than 2° linages. Later stages show low levels of expression in surrounding non-vulval hypodermal cells.

Around the time of the first VPC cell division, GFP expression from the *rgl-1* transcriptional fusion increases in presumptive 1° cells relative to presumptive 2° cells The significance of this change is unclear, since many transcriptional fusions of vulval signaling genes change expression patterns around this time (11-13, 20, 57; Rasmussen *et al.,* re-submitted). Yet one interpretation is that increased expression of RGL-1 in presumptive 1° cells accounts for resistance to *rgl-1(RNAi)* of this putative 1°-promoting activity of RGL-1.

### Tagged endogenous RGL-1 is expressed uniformly throughout VPC development

To validate the RGL-1 transcriptional fusion, we used the self-excising cassette (SEC) approach (58) to tag the endogenous 5’ end of the endogenous *rgl-1* locus with sequences encoding mNeonGreen fluorescent protein (mNG, FP) and a 3xFlag epitope tag (Fig. S3). We observed mNG throughout vulval lineages, consistent with that observed with the *rgl-1* promoter::gfp transcriptional fusion transgene.

### RGL-1 performs opposing GEF-dependent and GEF-independent functions in VPC fate patterning

A list of *rgl-1* mutations and other genetic tools is shown (Fig. 4A, B; Table 1). The Million Mutation Project (MMP) described random sequence identification of a large collection of mutagenized *C. elegans* lines, demanding only that mutant animals be viable and fertile over many generations (59). We analyzed the sole non-synonymous mutation in *ral-1, gk628801,* which caused an R139H mutation. Arg-139 is conserved in all Ras family members in metazoans. Outcrossed *ral-1(gk628801)* single mutant vulvae were superficially wild type (N = 83). In the *let-60(n1046gf)* background, *ral-1(gk628801)* increased 1° induction (Fig. 4C), consistent with our previously published analysis using *ral-1(RNAi)* and deletion mutations (Fig. 2; 20). Thus, *ral-1(gk628801)* perturbs RAL-1 2°-promoting signaling but does not disrupt RAL-1 sufficiently to confer exocyst-associated phenotypes.

**Figure 4:**
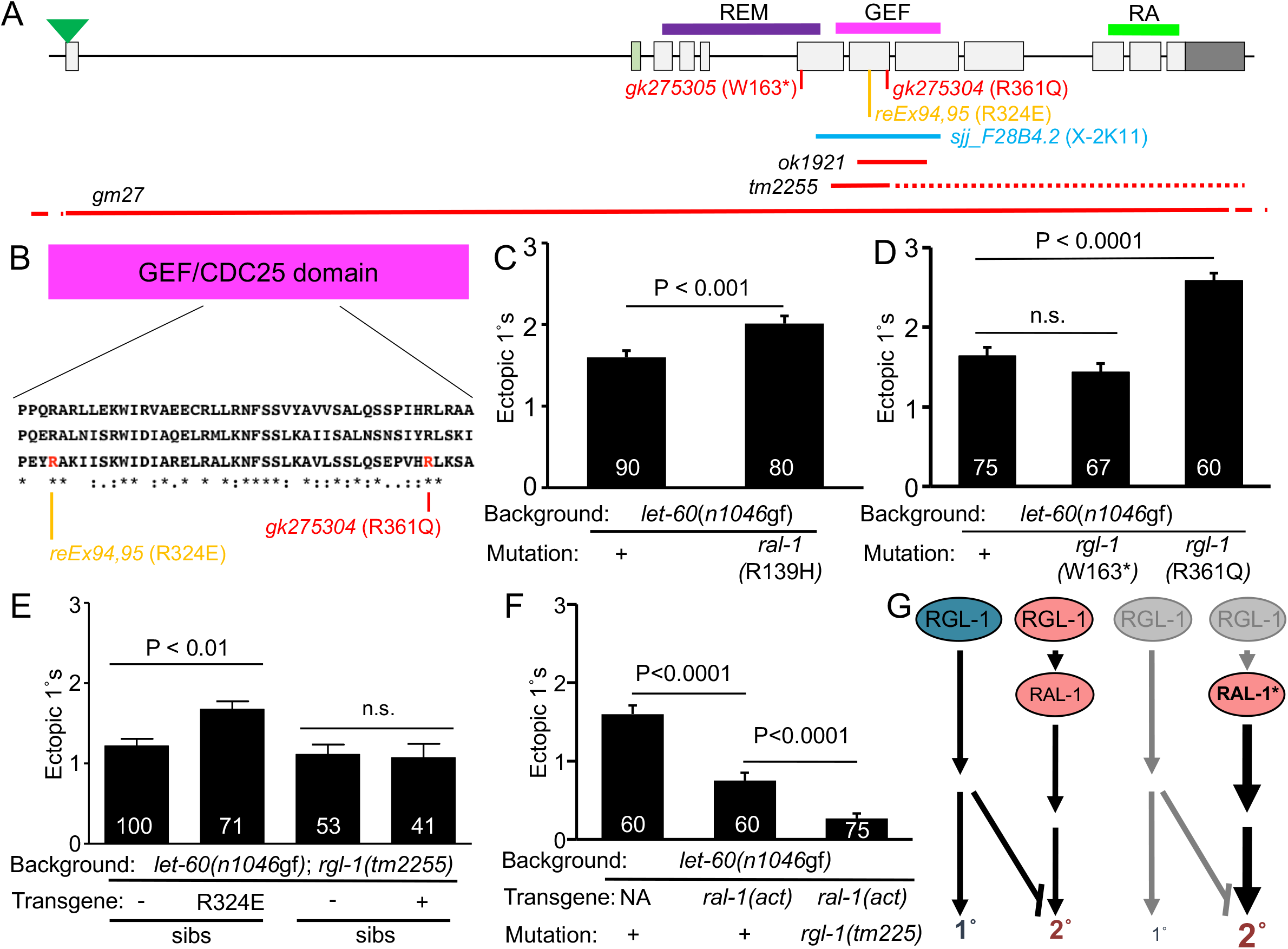
RGL-1 encodes genetically separable functions. **A)** A schematic of the *rgl-1* gene and genetic reagents for this analysis. Green triangle = mNeonGreen::3xFlag tag by CRISPR (Fig. S3). Light green exon 2: by RNAseq this is a rare mRNA species, and the 20 residues coded for by Exon 2 are not conserved among *Drosophila* and mammalian RalGEFs. Purple, pink and bright green lines: REM, GEF and RA (Ras Association) domains (The REM domain is a structural component of some but not all Ras GEFs). Red *gk* alleles: W163* and R361Q mutations from the million mutation project. Orange: canonical R324E “GEF dead “ mutation in transgenes *reEx94* and *reEx95.* Light blue: coverage of bacterially mediated RNAi clone (library location; (82)). Red lines: sequences deleted by *ok921, tm2255* and *gm27* deletions (dotted line indicates that deletion ends out of frame, and thus introduces premature stop codons). **B)** An alignment of a portion of the RalGEF domain containing the canonical “GEF dead” R324E and *gk275304* R361Q mutations (top to bottom: Human RGL2, in which the GEF dead mutation was validated, fly RalGEF, *C. elegans* RGL-1). **C)** The *ral-1(gk628801)* R139H missense mutation enhances 1 ° induction in the *let-60(n1046gf)* background. Data shown were scored concurrently and are representative of two assays. **D)** *rgl-1* mutation *gk275304* (R361Q) but not *gk275305* (W163*) enhances 1° induction in the *let-60(n1046gf)* background. Data shown were scored concurrently and are representative of two assays. **E)** Transgenic rescue of *rgl-1(tm2255)* in the *let-60(n1046gf)* background. Transgenic array-bearing animals harboring *Plin-31::rgl-1* cDNA with Pmyo-2::gfp co injection marker and their non-array-bearing siblings were scored. The *reEx94* (shown) and *reEx95* (not shown) R324E mutant transgenes enhanced 1° induction relative to their non-transgenic siblings, as scored in separate concurrent assays for each array, suggesting that a GEF-independent 1°-promoting activity of RGL-1 functions cell autonomously in the VPCs. *(reEx94 rgl-1* (R324E) mutant transgenic animals failed to respond to *ral-1(RNAi),* Fig. S4C.) The *reEx109* (shown) and *reEx110* (not shown) wild-type transgenes failed to alter 1° induction relative to their non-array-bearing siblings, as scored in separate concurrent assays for each array, suggesting that the GEF-dependent 2°-promoting activity of RGL-1 also functions cell autonomously in the VPCs. *(reEx109 rgl-1 (+)* transgenic animals responded to *ral-1(RNAi),* Fig. S4D.) **F)** *rels10[P_lin-31_::ral-1A(Q75L), P_myo-2_::gfp] (“ral-1(act)”)* suppressed ectopic 1° induction in the *let-60(n1046gf)* background; in a single concurrent assay, 1° induction was further suppressed by *rgl-1(tm2255),* revealing an opposing RGL-1 signal that may be 1°-promoting. Further validation of *reIs10 “ral-1(act)”* shown in Fig. S4B. Data are the mean ± standard error of the mean (SEM). For statistical reasons single, non-pooled assays are shown, and white numbers represent animals scored therein. Significance was calculated by Kruskal-Wallis, Dunn test. **G)** A schematic of the bypass experiment in F. Left: canonical and non-canonical RGL-1 activities are opposed and roughly equivalent (in the backgrounds assayed). Right: constitutive, VPC-specific activation of RAL-1 bypasses the GEF activity while also revealing a GEF-independent function of RGL-1, resulting in increased 2°-promoting signal and concomitant loss of 1°-promoting signal, with a net decrease of ectopic 1° cells.

Of 30 total non-synonymous mutations in *rgl-1,* we considered two likely to perturb function. *rgl-1(gk275304)* causes an R361Q change. Arg-361 is conserved in all CDC25/RasGEF domains. *rgl-1(gk275304)* conferred significant increase in ectopic 1° induction in the *n1046gf* background (Fig. 4D), consistent with disrupting GEF domain function and hence abrogating activation of RAL-1. *rgl-1(gk275305)* causes a W163* change, which did not alter ectopic 1° induction in the *let-60(n1046gf)* background. Both outcrossed single mutant strains were superficially wild type (N = 48 and 61, respectively). We therefore hypothesized that *rgl-1* encodes GEF-dependent and GEF-independent activities.

Using the VPC-specific *lin-31* promoter (60), in the *let-60(n1046gf); rgl-1(tm2255)* double mutant background we generated transgenic extrachromosomal arrays expressing VPC-specific *rgl-1(+)* or rgl-1(R324E), a mutation deficient in GEF catalytic activity (“GEF dead”) in mammalian RalGDS (RalGEF; 61). Ectopic 1° induction was scored in animals harboring extrachromosomal arrays, as indicated by pharyngeal GFP expressed by the *P_myo-2_::GFP* co-injection marker, vs. their siblings who had lost the transgene. Animals bearing the R324E transgenes showed significant increase in ectopic 1° induction. Animals bearing the wild-type transgenes were not different than their nonarray-bearing siblings (Fig. 4E). Consequently, we propose that RGL-1 performs GEF-dependent and GEF-independent functions that are genetically separable by mutating the GEF domain. We also conclude that the GEF-independent RGL-1activity functions cell autonomously in the VPCs.

Transgenic animals expressing VPC-specific RGL-1(+) should restore responsiveness to *ral-1(RNAi),* while transgenic animals expressing putative GEF-dead VPC-specific RGL-1(R324E) should not. We evaluated responsiveness of the *let-* 60(n1046gf); *rgl-1(tm2255)* background harboring each transgene. reEx94/95 transgenic R324E animals failed to respond to *ral-1(RNAi)* compared to control *gfp(RNAi)* (Fig. S4A; the 1° induction baseline is elevated due to rescue of the putative GEF-independent function shown in Fig. 4E). This observation was consistent with the absence of GEF activity. In contrast, animals harboring the *reEx109/110[P_lin-31_::rgl-1(+)]* transgene were responsive to *ral-1(RNAi),* resulting in increased 1° induction, consistent with VPC-specific rescue of GEF activity and thus cell autonomy of the GEF-dependent function of RGL-1 (Fig. S4B).

### Genetic bypass further reveals a GEF-independent activity of RGL-1

We previously showed that the ectopic vulva induction caused by mutation of *lin-31,* which confers ectopic 1° induction, was insensitive to perturbation of LET-60-LIN-45-MEK-2-MPK-1 1°-promoting signaling but sensitive to perturbation of LET-60-RGL-1-RAL-1 2°-promoting signaling, consistent with the model of LIN-31 functioning downstream of ERK/MAPK-1. In that assay, *rgl-1* function was assessed by RNAi, which resulted in increased 1° induction, as did depletion of *let-60* and *ral-1* (and *lin-12* positive control; 20). Here, *rgl-1(tm2255)* similarly increased ectopic 1° induction of *lin-31(301)* animals (Fig. S4C). This result is consistent genetically separable functions of RGL-1, and with the LIN-31/FoxB transcription factor functioning downstream of the putative non-canonical, GEF-independent signal, but in parallel to the canonical LET-60-RGL-1-RAL-1 signal (20).

We bypassed the requirement for GEF activity in RGL-1. We generated the *reIs10[P_lin-31_::ral-1A(Q75L), P_myo-2_::gfp]* integrated transgene (based on *reEx24* (20); control transgenes with RAL-1(+) had no effect). *reIs10* decreased ectopic 1° induction in the *let-60(n1046gf)* background (Fig. S4D). Into this background we additionally crossed the *rgl-1(tm2255)* out-of-frame deletion mutation and scored the three strains concurrently. *rgl-1(tm2255)* mutation significantly decreased ectopic 1° induction below the level observed with *reIs10* alone, nearly to wild-type levels (Fig. 4F). This result is consistent with further deletion of a 1°-promoting activity being revealed when RGL-1 is deleted and the GEF activity is bypassed by activated RAL-1. Taken together, these experiments suggest that *rgl-1* encodes two antagonistic signals, the GEF- and Ral-dependent 2°-promoting signal, and an unknown antagonistic signal, schematized in Fig. 4G.

### RGL-1 may function in the 1°-promoting PI3K-PDK-Akt cascade

Mammalian RalGDS binds to PDK and Akt1 in cultured cells, possibly functioning as a scaffold for PDK and Akt (45, 46). The AGE-1/PI3K-PDK-1 signal has been described as promoting 1° fate in VPC fate patterning (62), but AKT-1 was not implicated downstream of this process, potentially because of redundancy of AKT-1 and AKT-2 in *C. elegans* (63, 64). Previously, a gain-of-function mutation in AKT-1 was tested in a genetic background with reduced 1° induction: no effect was found, leading to the model that AKT-1 did not contribute to vulval induction (62). We re-evaluated the *akt-1(mg144gf)* mutation in the *let-60(n1046gf)* background and observed significant increase in ectopic 1 ° induction (Fig. 5A). To corroborate previously published results, we assessed the impact of both activated AKT-1 and PDK-1 in the hypo-induced *lin-45(n2506rf)* background: *akt-1(mg144gf)* did not alter the hypo-induced *lin-45(n2506rf)* phenotype, but *pdk-1(mg142gf)* did suppress the 1°-induction defect (Fig. S5C,D). This result, coupled with earlier analysis (62), suggests that the canonical AGE-1/PI3K-PDK-1/PDK-AKT-1/Akt cascade functions to promote 1° vulval fate. We speculate that hypo-induced presumptive 1° cells can respond to constitutively activated PDK-1 but not AKT-1, while in hyper-induced VPCs constitutively activated AKT-1 is sufficient to induce additional excess 1° cells. Perhaps this distinction between PDK-1 and AKT-1 reflects the presence of AKT-2, for which we do not have activating mutations.

**Figure 5.**
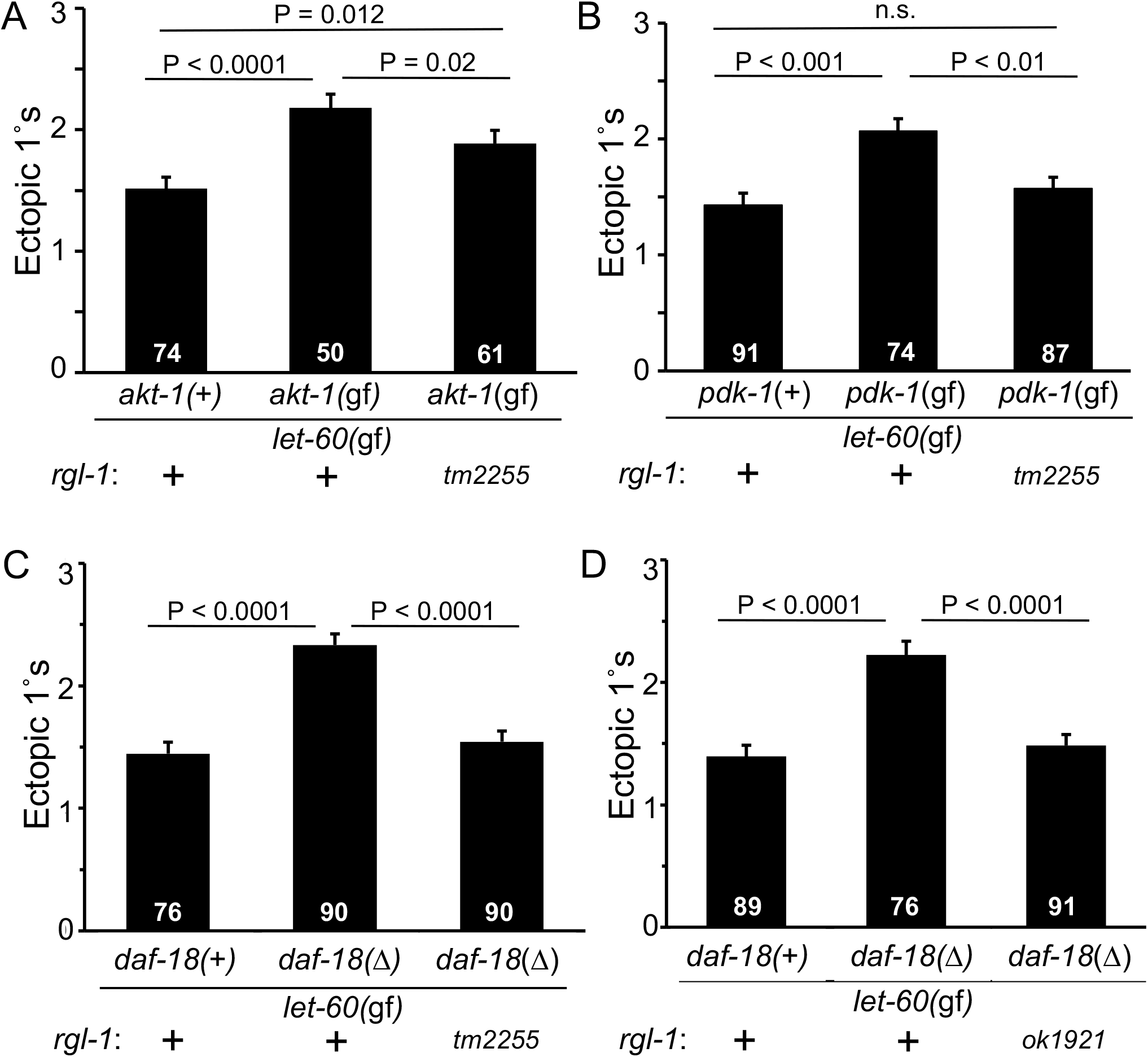
RGL-1 interacts genetically with the 1°-promoting AGE-1/PI3K-PDK-1-AKT-1 cascade. **A)** The constitutively activating *akt-1(mg144gf)* mutation increased promotion of 1 ° fate in the *let-60(n1046gf)* background, and this effect was partially blocked by *rgl-1(tm2255),* though the triple mutant was not suppressed to the double mutant baseline level. Animals were scored concurrently. These results were replicated with a re-built strain (Fig. S5A) and the strain with *rgl-1(ok1921)* (Fig. S5B). Data are the mean ± standard error of the mean (SEM). For statistical reasons single, non-pooled assays are shown, and white numbers represent animals scored therein. Significance was calculated by Kruskal-Wallis, Dunn test. **B)** The constitutively activating *pdk-1(mg142gf)* mutation increased promotion of 1° fate in the *let-60(n1046gf)* background, and was completely suppressed to baseline level by *rgl-1(tm2255).* Animals were scored concurrently and are representative of two assays. **C, D)** Mutation of the negative regulatory PTEN ortholog by *daf-18(ok480)* similarly increased 1° induction, and was completely suppressed to baseline level by *rgl-1(tm2255)* **(C)** and *rgl-1(ok1921)* **(D)**. Animals for each were scored concurrently and scoring was repeated once, with the same general results. *rgl-1(gk275305)* (nonsense; Fig S5C) but not *rgl-1(gk265304)* (GEF; Fig S5D) suppressed this *ok480* enhancement, suggesting that the pertinent RGL-1 activity is GEF-independent/non-canonical.

Including the *rgl-1(tm2255)* mutation in the *let-60(n1046gf); akt-1(mg144gf)* background significantly suppressed ectopic 1° induction, but not to the baseline of the *n1046gf* single mutant (Fig. 5A). Since we observed intermediate suppression, we constructed this strain twice and observed a similar result (Fig. S5A). We also reproduced this result with *rgl-1(ok1921)* and observed similar intermediate strength suppression that remained significantly above the baseline of the *let-60(n1046gf)* single mutant (Fig. S5B).

We further tested the relationship of RGL-1 with the rest of the PI3K cascade. Mutational activation of PDK-1 via the *pdk-1(mg142gf)* mutation also increased ectopic 1° induction in the *let-60(n1046gf)* background. This effect was completely suppressed by *rgl-1(tm2255)* (Fig. 5B), suggesting a quantitatively detectable difference between the epistatic relationships of PDK-1 and AKT-1 with RGL-1. Genetic disruption of DAF-18/PTEN, a negative regulator of this cascade in *C. elegans* in general (65, 66) and VPC fate patterning in particular (62), increases ectopic 1° induction in the *n1046gf* background. We found that *rgl-1(tm2255)* completely blocked this effect (Fig. 5B). The nonsense *rgl-1(gk275305)* but not the putative GEF dead *rgl-1(gk275305)* mutation similarly suppressed (Fig. S5E,F), suggesting that the putative RGL-1 1°-promoting is GEF-independent.

While mammalian RalGDS bound directly to PDK, RalGDS bound indirectly to Akt through the intermediary scaffold, JIP (JNK Interacting Protein; (45, 46)). Deletion the sole *C. elegans* JIP ortholog, JIP-1 (Fig. S5I) or RNAi depletion of JIP-1 (Fig. S5G) suppressed the ectopic 1 ° phenotype of the *daf-18(ok480) let-60(n1046gf)* double mutant, consistent with JIP-1 collaborating with RGL-1 to scaffold PDK-1 and AKT-1 1°-promoting signaling. RNAi depletion of both *pdk-1* and *rgl-1* similarly suppressed the enhanced 1° induction of *daf-18(ok480) let-60(n1046gf)* (Fig. S5H). However, the *jip-1* deletion allele enhanced *n1046gf* alone while suppressing *daf-18(ok480) n1046.* (Fig. S5I). We speculate that JIP-1 functions in the PI3K cascade, but may also function elsewhere in VPC fate patterning, perhaps in its canonical role as a scaffold for JKK and JNK MAP kinases. Further investigation of the role of JIP-1 is beyond the scope of this study.

A common target of the *C. elegans* PI3K-Akt cascade is inhibition of the DAF-16/FoxO transcription factor (67, 68). We tested the role of *daf-16* alleles *mu26, mu86* and *mgDf47* in different backgrounds, with inconclusive results. Consequently, we were unable to determine the role, if any, of DAF-16/FoxO functions in VPC fate patterning.

Taken together, these genetic epistasis experiments, using an assay of parallelism with *let-60(n1046gf),* suggest that RGL-1 contributes to the AGE-1/PI3K-PDK-1/PDK-AKT-1/Akt cascade, including the negative regulator lipid phosphatase, DAF-18/PTEN, in VPC fate patterning. Alone among the genetic tools used, the gain-of-function mutation in the downstream AKT-1 was only partially suppressed by *rgl-1(tm2255),* while the effect of other Akt cascade activators was completely suppressed by *rgl-1(tm2255).* We propose that RGL-1 is required for PDK-1 1°-promoting signal (and AGE-/PI3K, through inference from our DAF-18/PTEN deletion experiments), but only partially required for AKT-1 1° promoting signal. We note that mammalian Akt is activated via parallel mechanisms: phosphorylation by upstream PDK and binding of PIP3, resulting in recruitment to the plasma membrane and activation (69). If RGL-1 functions as a scaffold for PDK-1 and AKT-1, its deletion would be expected to result in reduction of the PDK-1 phosphorylation of AKT-1 but not PIP3-dependent recruitment of AKT-1 to the plasma membrane. Thus, a parsimonious interpretation of our data is that RGL-1 functions as a scaffold for PDK-1-AKT-1 signaling in 1° fate induction, and that this activity is independent of the GEF-dependent role of RGL-1 in promoting 2° fate through RAL-1 activation. However, we cannot rule out the possibility of parallelism between a GEF-independent RGL-1 activity and the AKT-1 cascade, or RGL-1 functioning in a bifurcated cascade downstream of AKT-1 that is not revealed by the mechanism of constitutively activate PDK-1 or DAF-18/PTEN upstream.

### Deletion of RGL-1 decreases fidelity of VPC patterning

Based on a relatively small sample size, RGL-1 deletions cause no gross VPC patterning defects, consistent with both cascades being modulatory, not central. We hypothesized that RGL-1 orchestrates activation of these two potentially opposing cascades – Akt output to presumptive 1 ° cells and Ral output to presumptive 2° cells – to improve robustness of the VPC developmental system in response to environmental stressors. A previous study investigated the impact on VPC fate patterning of environmental stressors in weakly disrupting genetic backgrounds, with an N2 baseline error rate of 0.2% under laboratory conditions (26). We similarly investigated the impact of environmental stressors of starvation, heat, and osmotic stress, compared to non-stressful conditions, on wild-type, *rgl-1(ok1921),* and *rgl-1(tm2255)* animals.

We evaluated 300 animals per genotype under each condition, totaling 1,200 animals per genotype and 3,600 animals overall (Fig 6A). To our surprise, the three measured environmental insults, compared to the unstressed, caused no patterning errors in either mutant or the wild type. However, the baseline patterning error rate was increased 15-fold in *rgl-1* mutants relative to the wild type, from 0.2% to 3.0%, with no difference between the stressed and unstressed animals (Fig. 6A). These results suggest a generalized increase in error rate in the absence of functional RGL-1. Consequently, we propose that RGL-1 does not impact robustness in response to environmental stressors. Rather, we hypothesize that RGL-1 mitigates patterning error within the complex signaling system that regulates VPC patterning. This observation represents intriguing and unprecedented insight into the role of signaling networks in developmental fidelity.

**Figure 6.**
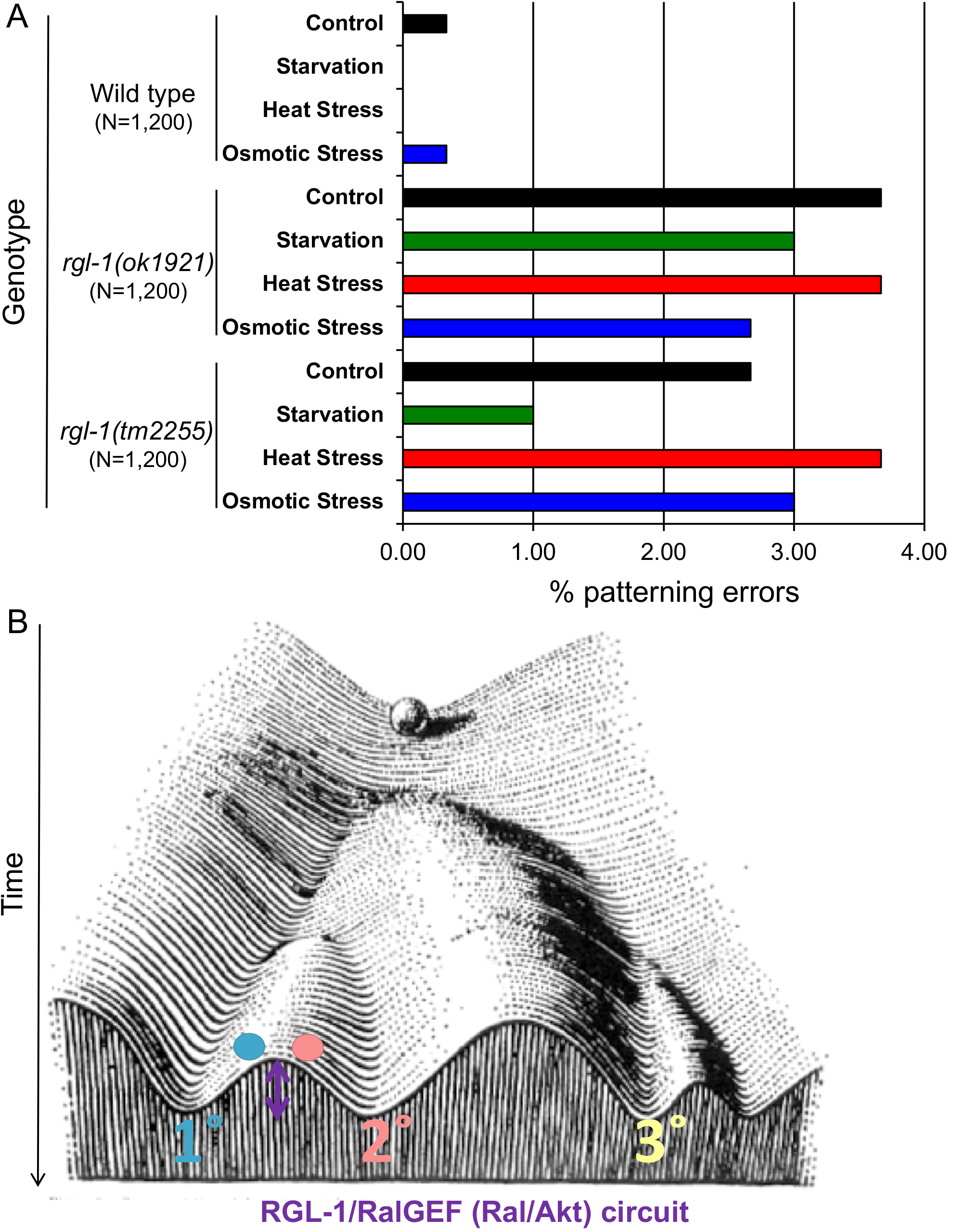
Deletion of RGL-1 increases patterning errors but not susceptibility to environmental stress. **A)** *rgl-1* deletions *ok1921* and *tm2255,* introgressed to the wild type for eight generations (DV2696, DV2697), caused increased patterning defects compared to the wild type. (N = 1,200 per genotype, pooled from 300 for each condition.) Yet no difference was observed between animals subjected to stressors vs. control animals. P < 0.0001 for *ok1921* or *tm2255* animals vs. the wild type, calculated by Kruskal-Wallis, Dunn test. **B)** A model for the role of Balanced Switches in VPC fate patterning, in the context of Waddington’s Developmental Topology. We hypothesize that through decreasing error rate, the RGL-1 signaling circuit effectively increases the “barrier” between 1° and 2° fates.

## DISCUSSION

The impetus for this analysis was two enigmatic genetic observations. First, deletion of RGL-1 confers no overt phenotype, in contrast to deletion of its signaling partner, RAL-1. Second, deletion of RGL-1 confers no net effect on 1° vs. 2° induction, while deletion or depletion of RAL-1 specifically reduces 2° induction. Thus, in two different ways RGL-1 is non-equivalent to RAL-1. By pursuing these two genetic observations, we arrived at important models regarding the roles of RalGEF and Ral in signal transduction and development.

### RAL-1 but not RAL-1 is essential for development

RAL-1 was previously implicated in developmental functions. Yet despite the ostensible linearity of Ras-RalGEF-Ral signaling, we unexpectedly found that RGL-1 is inessential for development. No role for LET-60/Ras has been found in exocyst and PAR complex function (70). Based on our previous analysis of the *C. elegans* RalGAP, neither do mutations in the HGAP-1/2 (54), predicted to confer excess activation of endogenous RAL-1. Nor does a constitutively active RAL-1 generated by CRISPR introduction of the G26V mutation into the endogenous *ral-1* gene (Shin, *et al*., in press). Given that deletion of neither the GEF nor the GAP disrupts exocyst-dependent developmental events, but deletion of the small GTPase itself, RAL-1, does so, we hypothesize that the RAL-1 role as a membrane tether for the exocyst is independent of nucleotide-bound state. This hypothesis does not imply that there is no activation-dependent alteration of exocyst function by RAL-1, merely that such is not essential for the described developmental events.

An alternative hypothesis is that other GEFs or GAPs function redundantly with RGL-1 and HGAP-1/2 to regulate the GDP/GTP cycle of RAL-1. Based on our analysis, RGL-1 is required for GTP loading on RAL-1 for the 2°-promoting signal. But this other GEF would need to be specific for RAL-1 association with the exocyst and their functions in development, but not signaling to promote 2° fate. While we cannot exclude this possibility, no GEFs fitting that role have been described in any system (RalGPS, a mammalian Ral-selective GEF that contains SH3 and PH domains, is not encoded in the *C. elegans* genome; (20), nor is TD-60/RCC2, similar to the Ran GEF RCC1 that controls nuclear import/export, which has been proposed to be an atypical GEF for RalA (71)).

### RGL-1 performs genetically separable and opposing functions in VPC fate patterning

To our surprise, we found that deletion alleles of *rgl-1* caused no net alteration of the balance of 1° and 2° VPCs in the *let-60(gf)* background. By genetic analysis, RGL-1 performs its canonical role in promoting 2° fate as a signaling intermediary between LET-60/Ras and RAL-1 ((20); this study), a role that mirrors extensive biochemical and cell biological evidence from mammalian cell culture (reviewed in 35, 42, 72). Here, further genetic analysis reveals an additional, unexpected role for RGL-1, that of a non-canonical signaling participant opposing the canonical role as an activator of RAL-1. This non-canonical role apparently counteracts or “cancels out” the canonical role. In sensitized backgrounds, putative null mutations have no net effect on the sensitive balance between 1° and 2° VPC fates. We do not intend to imply that opposing functions are “equal,” only that in most of our assays they appear to counter-balance each other with approximate equivalency. The observation that deletion of RGL-1 increase the basal error rate 15-fold supports the idea that RGL-1 serves an important function, that of fidelity.

The key biochemical connection between RalGEF, PDK, and Akt came in mammalian cell culture, and in that context RalGEF-Ral and PDK-Akt cascades worked in concert (45). Yet historically RalGEF-Ral signaling has also been found to oppose canonical Ras effectors in cell culture (73). We hypothesize that the relationship between these cascades may depend on cell context, and in the case of vertebrates, which paralog of RalGEF is expressed.

RGL-1 is not the first protein in the VPC fate patterning network found to be bifunctional and promote both 1° and 2° fates. Depending on signaling dose and via mechanisms we do not understand, LIN-3/EGF-LET-23/EGFR signaling was found to promote both 1° and 2° fates, resulting in the original Morphogen Gradient Model (1, 2, 4). We found that LET-60/Ras can promote 1° or 2° fate depending on its use of effector, LIN-45/Raf or RGL-1/RalGEF (19, 20). Yet the situations are not equivalent. LIN-3, LET-23, and LET-60 are essential for vulval induction: strong loss results in complete absence of 1° fate induction and hence a vulvaless phenotype (74–77), and thus their roles in 2° fate induction were teased out only in sensitized backgrounds or special assays (1, 2, 20). In contrast, RGL-1 is dispensable for 1° and 2° fate induction, permitting dissection of its balanced and opposing functions. Furthermore, deletion of RGL-1 does not alter any other known developmental events.

These observations positioned us to discover an unexpected aspect of RGL-1 function: it contributes to two modulatory cascades, neither of which is essential for VPC induction. And since the two cascades promote opposing outcomes, loss of both together has no net effect on the delicate balance between 1° and 2° fates. Consequently, this unusual feature of RGL-1 function in vulval signaling, in the anatomically simple nematode that mostly lacks paralog redundancy, provided the opportunity to discover what may be a heretofore unknown property of signaling networks. We speculate that we have identified a signaling switch that is dispensable for development, and that precise regulation of this switch increases the fidelity of signaling networks and hence development. Specifically, in this case RGL-1 may orchestrate the opposing outputs of AKT-1, which promotes 1° fate through unknown downstream targets (this study), and RAL-1, which promotes 2° fate through EXOC-8/Exo84 of the exocyst, the GCK-2/MAP4 kinase, MLK-1/MAP3K, and PMK-1/p38 MAP kinase (Shin et al., in press).

### Linking opposed signaling cascades and mitigating development noise

The problem that needs to be “solved” by the VPC fate patterning system is how to strictly define developmental fields in response to an initial point source of LIN-3/EGF ligand. In other words, how does one generate the precise 3°-3°-2°-1°-2°-3° pattern with 99.8% accuracy without mis-specified or ambiguous fates that might block mating and egg laying? A critical question, then, is how the programming of signal transduction networks decreases the potential for errors, or developmental “noise.” Previously, the fidelity of VPC fate patterning was thought to be a property that emerges from the combination of three mechanisms: 1) Sequential Induction sets up the basic pattern, 2) the Morphogen Gradient collaborates with Sequential Induction to more precisely sculpt the initial pattern, and 3) Mutual Antagonism serves to exclude potentially conflicting signals from cells that are initially specified, thus preventing assumption of wrong or ambiguous fates. We speculate that the roles we describe here for RGL-1 define a fourth method that is woven into the other three: orchestration of two modulatory cascades to more sharply demarcate fates as a function of the VPC’s spatial relationship to the AC and other VPCs. We name this property “Balanced Switches.”

Yet is the participation of RGL-1 in two opposing cascades necessarily a novel mechanism that promotes fidelity? Perhaps not. Perhaps RGL-1 as a point of intersection between two conserved but opposing cascades is coincidental. In this alternative model, the loss of fidelity observed in deletion mutants of *rgl-1* may be merely the consequence of losing two independent modulatory cascades, with AKT-1 and RAL-1 outputs, that each reinforces their respective fates. In this model, the loss of fidelity observed in *rgl-1* deletion mutants is happenstance, a side effect of losing roughly equal and opposite modulatory cascades.

Alternatively, the wiring of these two cascades together in opposition could be of mechanistic significance. How such a mechanism would function is unclear. One possibility is that the RGL-1 “Balanced Switch” functions as an “Insulated Switch.” The simplest way to envision this model is through mechanisms of subcellular recruitment and sequestration: RGL-1 engagement in one signal could physically exclude its engagement in the opposing signal, perhaps because the RGL-1 protein has been re-localized to a portion of the cell insufficient to participate in both signals, or is concomitantly modified to prevent interaction in its complementary function. For example, binding of Ras to the RA domain presumably recruits RGL-1 to the plasma membrane, consistent with Ras interactions with other effectors in mammalian cells. This recruitment could sequester RGL-1 away from the subcellular compartment in which PDK-1 and AKT-1 signal. Conversely, RGL-1 bound as a scaffold to PDK-1 and AKT-1 could be sequestered away from the subcellular compartment in which Ras interacts with RGL-1, the preventing participation in the canonical LET-60-RGL-1-RAL-1 2°-promoting cascade. A metaphor for such a switch is the two-headed Pushmi-Pullyu from Dr. Doolittle: when one head pulls forward, the other is by necessity pulled back, and vice-versa. An inessential signaling protein with this property – like RGL-1 – could function to reinforce two different cell fates to improve developmental fidelity. And the mutually exclusive bi-directionality of such a switch would ensure that inappropriate signaling is minimized.

The mechanism by which RGL-1 contributes to different fates remains to be determined. We do not yet have biomarkers for RAL-1 or AKT-1 output in the VPCs. And because of the brief developmental window of VPC patterning and the tiny volume of VPC lineages relative to the entire animal, biochemical approaches are inadequate. The concept of the “Balanced Switch” being woven into signaling networks is a fascinating one, and one difficult to test with the typical manifold gene redundancy present in mammalian systems, or the essential nature of many signaling genes in the developmentally more complex *Drosophila.* Perhaps developmental patterning of the *C. elegans* vulva is the right place to test this idea, but more tools are needed to do so. Yet, in the dawn of the CRISPR era, we remain confident about our ability to do so in the future, including the possibility of generating two *rgl-1* genes, each of which governs one by not the other function, *i.e.* uncoupling the two halves of the “switch.”

### Developmental fidelity vs. developmental robustness

Increasing noise could impact two obvious outcomes: fidelity and robustness. Fidelity here we define as developmental accuracy in the absence of external environmental perturbation, while robustness we define as developmental accuracy in the presence of perturbation. To our surprise, deletion of RGL-1 altered the fidelity but not robustness of VPC development: environmental perturbations had no effects on wild-type vs *rgl-1* deletion mutant animals, while all mutant animals had a 15-fold increase in patterning error rate compared to the wild type, regardless of their environment. Conversely, small changes in function, from weak mutation of certain cascades or introgression into different genetic backgrounds that might harbor mutations, caused increased sensitivity to environmental insult (26, 30). Yet, surprisingly, a four-fold change in EGF dose did not appreciably alter VPC patterning (27). We speculate that it is *coordinate* change caused by deletion of RGL-1 that does not decrease robustness. In other words, perhaps it is disruption of the delicate balance of 1°- and 2°-promoting signals that sensitizes the system to environmental insult. Yet accumulation of mutations over many (~250) generations does increase error rate (28), and thus in these background mutations may be identified that disrupt fidelity-promoting functions like that of RGL-1.

### Conclusions

We define a putative regulatory system, which we term “Balanced Switching,” that potentially mitigates potential signaling noise and hence developmental error. This system would likely not be discovered in experimental platforms with extensive paralog redundancy, or where the genes in question are essential for viability or the process being studied. Whether Balanced Switches are generalizable to other systems, and the molecular details by which they function, await further analysis.

In addition to divergent functions of RGL-1 in signaling, we also delineate mechanistic details of RAL-1 vs. RGL-1 function in essential functions performed by the exocyst and PAR complexes. Our results suggest that the role of RAL-1 in central machinery of cell biology is independent of GDP/GTP state, though we cannot rule out some switchable modulation of exocyst and PAR complex regulation by RAL-1.

## Supplemental Information

Supplemental information includes seven Supplemental figures.

### Methods

#### *C. elegans* handling and genetics

Nomenclature is as described (78, 79). All strains were derived from the N2 wild type. Except where noted, animals were cultured on NGM agar plates with OP50 bacteria on at 20°C (80). Strains used are shown in Table S1. Data were analyzed with GraphPad Prism software (GraphPad Software Inc., La Jolla, CA).

PCR primers are listed in Table S2. For single animal genotyping PCR we used Taq PCR Master Mix (Qiagen): every reaction was run concurrently with +/+, m/+ and m/m control animals. The genotype of every PCR-genotyped strain was double-checked after completion of the strain construction. For each newly analyzed mutation, PCR products were sequenced to confirm break points. *rgl-1(ok1921)* was detected by triplex PCR using primers DJR614/615/616 (Tm: 59°, 35 cycles), resulting in 366 bp (wild type) and 233 bp *(ok1921)* bands (Fig. S6). Point mutations in *rgl-1* were tracked in *trans* to *ok1921.* Early constructions detected *rgl-1(tm2255)* by triplex PCR using primers TZ20/DJR614/DJR615 (Tm: 58°, 35 cycles), resulting in 913 bp (wild-type) and 595 bp *(tm2255)* bands (Fig. S7). Later constructions detected *rgl-1(tm2255)* by triplex PCR using primers FSM7/8/9 (Tm: 58°, 35 cycles), resulting in 509 bp (wild-type) and 254 bp *(tm2255)* bands. *pdk-1(mg142gf)* was amplified by primers DRC1/2 (Tm: 60°, 35 cycles) to generate a 426 bp band, and digested with Hpa II (NEB; 5 units added to total reaction volume doubled with water with NEB buffer #1 to 0.5x total, digested overnight at 37°C). The band from the wild-type allele was digested to yield 126 and 300 bp bands, while the *mg142* lesion abolishes the Hpa II site. *daf-18(ok480)* was detected by triplex PCR using primers FSM4/5/6 (Tm: 54°, 35 cycles), resulting in 388 bp (wild-type) and 216 bp *(ok480)* bands. For this study, *dpy-9(e14)* was used as a balancer for *daf-18(ok480),* and the *ok480* genotype confirmed after construction. *akt-1(mg144gf)* was balanced during strain constructions by *dpy-11(e224) unc-76 (e905). ral-1(gk628801)* was detected by amplification with primers DJR778/779 (Tm = 59.9, 35 cycles) to generate a 250 bp band and digested with HpyCH4 IV (NEB; 5 units added to total reaction volume doubled with water with NEB buffer #1 to 0.5x total, digested overnight at 37°C). The band from the wild-type allele was digested to yield 122 and 128 bp bands, while the *gk628801* lesion abolishes the HypCH4 IV site.

Transgenic extrachromosomal array *reEx24[P_lin-31_::ral-1(Q75L), P_myo-2_::gfp)]* (20) was integrated by irradiation of late L4 animals using a Stratalinker (Stratagene) at dose of 12 mJ/cm^2^. 451 F2 progeny were screened for integration to obtain *rels10 [P_lin-31_::ral-1(Q75L), P_myo-2_::gfp)]. reIs10* was mapped to the region of Chromosome I +5.

#### VPC induction assays

VPC induction was analyzed on an agar pad (molten 3% NG agar with 5 mM sodium azide) on a slide. Late L4 animals were added to a 5 μl drop of M9 buffer and the cover slip added. Ectopic pseudovulvae were scored at 600x or 1000x as invaginations using DIC/Nomarski optics (Nikon eclipse Ni) with images captured using NIS-Elements AR 4.20.00 software. Ectopic 1° vulval induction index, from 0 to 3 ectopic 1°s, is described elsewhere ((20); Shin et al in press). Under-induced backgrounds were scored as total VPCs induced, typically 0-3 (in under-induced backgrounds, 2° lineages are occasionally missing). To summarize, we scored the wild-type vulval induction based on the stereotypical 2°-1°-2° lineages centered on the AC at the A-P midpoint of the gonadal primordium (forming the “Christmas T ree” or “Stanley Cup” shape). In the *let-60(n1046gf)* and *let-23(sa62gf)* backgrounds, the morphology of ectopic 1 ° cells generally conformed with the described “cap” structure where the entire 1° lineage has pulled away from the cuticle (2). We did not observe ectopic pseudovulvae with the characteristic asymmetrical “beret” lineage of 2° lineages, where one side remains attached to the cuticle.

As described previously ((20); Shin et al in press), we frequently observed drift of the severity of the *let-60(n1046gf)* but not *let-23(sa62gf)* Muv phenotype, generally increasing the severity of ectopic 1° induction. For this study we analyzed parental MT2124 and outcrossed strains harboring the *n1046gf* single mutant, and established that undrifted strains average 1.2-1.5 ectopic 1°s. (When grown on bacterially mediated RNAi food source HT115, induction was consistently higher; (20); Shin et al in press). For all strains harboring the *n1046gf* mutation we employed stringent scoring criteria: the *n1046gf* single mutant was scored first, and experiments deviating from the expected range of induction (1.2-1.5, 1.5-1.8 for RNAi) were discarded. We additionally always worked with freshly thawed or chunked strains, pulling from a large collection of undrafted strains that had been frozen. Thus, we always worked with animals that were freshly derived from a cross or from a thaw. When using this rigorous protocol, we rarely observe significant deviations from expected baselines. Additionally, we only compared genotypes that were scored concurrently, thus minimizing variation from assay to assay.

### Bacterially mediated RNA interference

RNAi was performed as described previously ((20); Shin in press). RNAi plasmids used were: pREW2 *(luciferase/luc;* Shin *et al.* in press), X-2K11 *(rgl-1),* III-7M13 *(ral-1),* I-1K04 *(pop-1),* and *gfp* (20). Each RNAi clone was sequence verified. HT115 bacteria were used as the host for RNAi clones (81). Bacteria were grown on NGM plates supplemented with 50 μg/ml carbenicillin and 1 mM IPTG. Bacteria were grown (but not overgrown) overnight, without antibiotic selection. 80 μl of fresh culture was seeded on plates on day 1, grown overnight, L4 animals were added on day 2, transferred to a fresh plate on day 3, and scored on day 5.

### Plasmid subcloning and transgene generation

Using primers RGL-1F and RGL-1R, the *rgl-1a* cDNA was amplified from clone yk643d11, and digested with Bgl II and Not I. This isoform lacks exon 2, which by RNAseq data is rare (Wormbase WS263), yet still rescues, arguing that exon-2 is not required for vulval signaling. Plasmid vector pB255, which contains the *lin-31* promoter and additional regulatory sequences and drives expression in VPCs (60), was digested with Bgl II and Not I to receive the *rgl-1* insert. The putative R324E “GEF dead” mutation was introduced by PCR with Pfu Turbo using primers KM1 and KM2. The resulting plasmids, *P_lin-31_::rgl-1(+)* and *P_lin-31_::rgl-1(R324E),* were injected at 5 ng/μl along with co-injection marker pPD118.33(P_myo-2_::gfp) at 5 ng/μl into strain DV2190 *let-60(n1046gf); rgl-1(tm2255)* to generate arrays *reEx109* and *reEx110 (rgl-1(+))* and *reEx94* and *reEx95 (rgl-1(R324E)),* which express wild-type and “GEF dead” RGL-1, respectively, specifically in VPCs.

### CRISPR/Cas9-dependent genome editing

*rgl-1(re179[mNeonGreen::3xFlag::rgl-1])* was generated using the positive-negative selection self-excising cassette method (58). The repair template for *rgl-1* 5’ tagging was generated by Gibson Assembly (NEB) with digested target SEC vector pDD268, and ~500 bp of homology arms amplified from genomic DNA by Q5 polymerase (NEB). We used two sgRNA sequences: (#1) 5’-ACACCTTCGTATCCTTGTGG<u>CGG</u>-3’ and (#2) 5’-GGTCTGAGTTCTTCTGACGA<u>TGG</u>-3’, and hence generated two targeting vectors and one repair template. mNG^^^3xFlag::RGL-1 was generated by microinjection of the repair template (20 ng/μl), sgRNA-Cas9 #1 (25 ng/μl), sgRNA-Cas9 #2 (25 ng/μl), and injection marker *P_myo-2_::mCherry* (2.5 ng/μl) into wild-type animals. Knock-in alleles Genotyping and sequencing of *rgl-1* 5’ CRISPR tagging was performed with HS125/126/127 (Tm=54). Validation was performed by western blotting using monoclonal anti-Flag antibody (Sigma-Aldrich F1804) (1:2000), monoclonal anti-α-tubulin antibody (Sigma-Aldrich T6199) (1:2000) and goat anti-mouse secondary antibody (MilliporeSigma 12-349) (1:5000).

For reasons unknown, all seven *rgl-1* CRISPR alleles generated harbored mutations. Repair templates were re-checked by sequencing to confirm that sequences were wild type. DV3225 *rgl-1(re179[mNeonGreen::3xFlag::rgl-1])* harbored only promoter mutations (C insertion at -614, C deleted at -375, C to T at -395) and so was selected for further analysis. Analysis of the ModEncode database showed no peaks of promoter occupancy at these sites. *rgl-1(re179[mNeonGreen::3xFlag::rgl-1])* had no effect on 1° induction in the *let-60(n1046gf)* background (P = 0.57 between strains with and without the *re179* insertion, N = 86 and 78, respectively).

### DIC, epifluorescence and confocal microscopy

For DIC scoring of induction, live and appropriately staged animals were mounted in 5 mM sodium azide/M9 buffer on slides with 3% agar pad. For epifluorescent imaging, animals were mounted in 2 mg/ml tetramisole/M9 buffer and visualized using a Nikon Eclipse TE2000U microscope equipped with a DVC-1412 CCD camera (Digital Video Camera Company), with Hamamatsu SimplePCI acquisition software. Confocal images were captured by A1si Confocal Laser Microscope (Nikon) using NIS Elements Advanced Research, Version 4.40 software (Nikon).

### Assessment of patterning error rate with environmental insults

Large-scale vulval induction and environmental stressors were scored as previously described by Braendle *et al.* (26).

## Acknowledgements

We thank Y. Kohara for the *rgl-1* cDNA clone, the National Bioresource Project for the Experimental Animal *Caenorhabditis elegans* and the *C. elegans* Gene Knockout Consortium (Mitani, Barstead and Moerman labs) for deletion strains, D. Baillie for BC14985, and the *Caenorhabditis* Genetics Center (CGC), which is funded by NIH Office of Research Infrastructure Programs (P40 0D010440). We thank A. Fanning, B. Goldstein and K. Caron at UNC for microscope use and L. Vergara and the Center for Advanced Imaging at the TAMU Institute of Biosciences and Technology in Houston. We thank the Reiner lab for helpful discussions. This work was supported by NIH grant GM121625 to D.J.R.

---

**Supplementary Table 1.** Strains

| Strain | Genotype | Application |
| --- | --- | --- |
| DV3311 | *ral-1(tm2760) / qC1 [dpy-19(e1259ts) glp-1(q339) nls189[P<sub>myo-2</sub>::gfp]]* | Inspecting mutant VPCs |
| DV2924 | *ral-1(tm5205) / qC1 [dpy-19(e1259ts) glp-1(q339) nls189[P<sub>myo-2</sub>::gfp]]* | Inspecting mutant VPCs |
| MT2124 | *let-60(n1046gf) IV* | Fig. 2B,C,E, baselines throughout |
| DV2214 | *let-60(n1046gf) IV (2x outcrossed)* | Results |
| DV2215 | *let-60(n1046gf) IV (2x outcrossed)* | Results |
| RB1576 | *rgl-1(ok1921) X* | Starting reagent |
| DV2194 | *rgl-1(ok1921) X (4x outcrossed)* | throughout |
| FX2255 | *rgl-1(tm2255) X* | Starting reagent |
| DV2175 | *rgl-1(tm2255) X (4x outcrossed)* | throughout |
| DV2937 | *ral-1(tm5205) III / qC1 [dpy-19(e1259ts) glp-1(q339) nls189[P<sub>myo-2</sub>::gfp]]; let-60(n1046gf) IV* | Fig. 2F |
| DV2190 | *let-60(n1046gf) IV; rgl-1(tm2255) X* | Fig. 2D,F,G |
| DV2248 | *let-60(n1046gf) IV; rgl-1(ok1921) X* | Fig. 2F,G |
| DV2251 | *let-60(n1046gf) IV; lon-2(e678) X* | Fig. 2G |
| DV2252 | *let-60(n1046gf) IV; fax-1(gm27) lon-2(e678) X* | Fig. 2G |
| DV2191 | *let-60(n1046gf) IV; arls92[Pegl-17::NLS-cfp::LacZ + Pttx-3::gfp] V* | Fig. S2A, C, D |
| DV2750 | *let-60(n1046gf) IV; arls92[Pegl-17::NLS-cfp::LacZ + Pttx-3::gfp] V; rgl-1(tm2255)* | Fig. S2B, E |
| WU49 | *lin-45(n2506rf) unc-24(e138) IV* | Fig. S2F, S5E, S5F |
| DV2783 | *lin-45(n2506rf) unc-24(e138) IV; rgl-1(tm2255) X* | Fig. S2F |
| PS1524 | *unc-4(e120) let-23(sa62gf) II* | Fig. S2G |
| DV2958 | *unc-4(e120) let-23(sa62gf) II; rgl-1(tm2255) X* | Fig. S2G |
| BC14985 | *dpy-5(e907) I; sEx14985[rCesF28B4.2::gfp + PCeh361(dpy-5(+))]* | Fig. 3 |
| DV3312 | *rgl-1(re179[mNeonGreen^3xFlag::rgl-1]) X* | Fig. S3 |
| VC40420 | *ral-1(gk628801rf) III (unoutcrossed)* | Results |
| DV2942 | *ral-1(gk628801rf) III (6x outcrossed)* | Results |
| DV2799 | *ral-1(gk628801) III; let-60(n1046gf) IV* | Fig. 4C |
| VC20011 | *rgl-1(gk275304) X (unoutcrossed)* | Results |
| DV2883 | *rgl-1(gk275304) X (4x outcrossed)* | Results |
| VC40052 | *rgl-1(gk275305) X (unoutcrossed)* | Results |
| DV2884 | *rgl-1(gk275305) X (4x outcrossed)* | Results |
| DV2764 | *let-60(n1046gf) IV; rgl-1(gk275304) X* | Fig. 4D |
| DV2765 | *let-60(n1046gf) IV; rgl-1(gk275305) X* | Fig. 4D |
| DV2537 | *let-60(n1046gf) IV; rgl-1(tm2255) X; reEx94[P<sub>lin-31</sub>::rgl-1(R324E) + P<sub>myo-2</sub>::GFP]* | Fig. 4E, S4A |
| DV2538 | *let-60(n1046gf) IV; rgl-1(tm2255) X; reEx95[P<sub>lin-31</sub>::rgl-1(R324E) + P<sub>myo-2</sub>::GFP]* | Fig. 4E, S4A |
| DV2736 | *let-60(n1046gf) IV; rgl-1(tm2255) X; reEx109[P<sub>lin-31</sub>::rgl-1(+) + P<sub>myo-2</sub>::GFP]* | Fig. 4E, S4B |
| DV2737 | *let-60(n1046gf) IV; rgl-1(tm2255) X; reEx110[P<sub>lin-31::rgl-1(+)</sub> + P<sub>myo-2::GFP</sub>]* | Fig. 4E, S4B |
| MT301 | *lin-31(n301) II* | Fig. S4C |
| DV2763 | *lin-31(n301) II; rgl-1(tm2255)* | Fig. S4C |
| DV2140 | *let-60(n1046gf) IV; reEx24[P<sub>lin-31::ral-1(Q75L)</sub>, P<sub>myo-2::gfp</sub>]* | Zand, 2011, derived *rels10* |
| DV2335 | *rels10[P<sub>lin-31::ral-1(Q75L)</sub> + P<sub>myo-2::GFP</sub>] I* | Results |
| DV2699 | *rels10[P<sub>lin-31::ral-1(Q75L)</sub> + P<sub>myo-2::GFP</sub>] I; let-60(n1046gf) IV; let-60(n1046gf) IV* | Fig. 4F, S4D |
| DV2700 | *rels10[P<sub>lin-31::ral-1(Q75L)</sub> + P<sub>myo-2::GFP</sub>] I; let-60(n1046gf) IV; let-60(n1046gf) IV; rgl-1(tm2255) X* | Fig. 4F |
| CB2065 | *dpy-11(e224) unc-76(e905)* | *akt-1* balancer |
| GR1310 | *akt-1(mg144gf) V* | Fig. 5 |
| DV2746 | *let-60(n1046gf) IV; akt-1(mg144gf) V* | Fig. 5A, S5A |
| DV2747 | *let-60(n1046gf) IV; akt-1(mg144gf) V; rgl-1(tm2255) X* | Fig. 5A, S5A |
| GR1318 | *pdk-1(mg142gf) X* | Fig. 5 |
| DV2773 | *let-60(n1046gf) IV; pdk-1(mg142gf) X* | Fig. 5B, S5B |
| DV2782 | *let-60(n1046gf) IV; pdk-1(mg142gf) X* | Fig. 5B |
| DV2844 | *let-60(n1046gf) IV; pdk-1(mg142gf) X* | Fig. 5B |
| SP934 | *unc-1(e538) dpy-3(e27) X* | Fig. 5B |
| DV2774 | *unc-1(e538) dpy-3(e27) rgl-1(tm2255) X* | Fig. 5B |
| DV2819 | *pdk-1(mg142gf) rgl-1(tm2255) X* | Fig. 5B |
| DV2822 | *let-60(n1046gf) IV; pdk-1(mg142gf) rgl-1(tm2255) X* | Fig. 5B |
| DV2791 | *lin-45(n2506rf) unc-24(e138) IV; akt-1(mg144gf) V* | Fig. S5F |
| DV2795 | *lin-45(n2506rf) unc-24(e138) IV; pdk-1(mg142gf) X* | Fig. S5E |
| RB712 | *daf-18(ok480) IV* | Starting reagent |
| DV2855 | *daf-18(ok480) 2x outcrossed* | Fig. 5 |
| DV2510 | *daf-18(ok480) let-60(n1046gf) IV* | Fig. 5C, D, S5C, S5D, S5G, S5H |
| DV2535 | *daf-18(ok480) let-60(n1046gf) IV; rgl-1(tm2255) X* | Fig. 5C |
| DV2509 | *daf-18(ok480) let-60(n1046gf) IV; rgl-1(tm2255) X* | Fig. 5D |
| DV2679 | *jip-1(km18) II* | Fig. S5I |
| DV3013 | *jip-1(km18) II; let-60(n1046gf) IV* | Fig. S5I |
| DV3358 | *jip-1(tm6137) II (4x outcrossed)* | Fig. S5I |
| DV3505 | *jip-1(tm6137) II; let-60(n1046gf) IV* | Fig. S5I |
| DV3506 | *jip-1(tm6137) II; daf-18(ok480) let-60(n1046gf) IV* | Fig. S5I |
| DV2696 | *rgl-1(ok1921) 8x outcrossed* | Fig. 6 |
| DV2697 | *rgl-1(tm2255) 8x outcrossed* | Fig. 6 |

**Supplementary Table 2.**
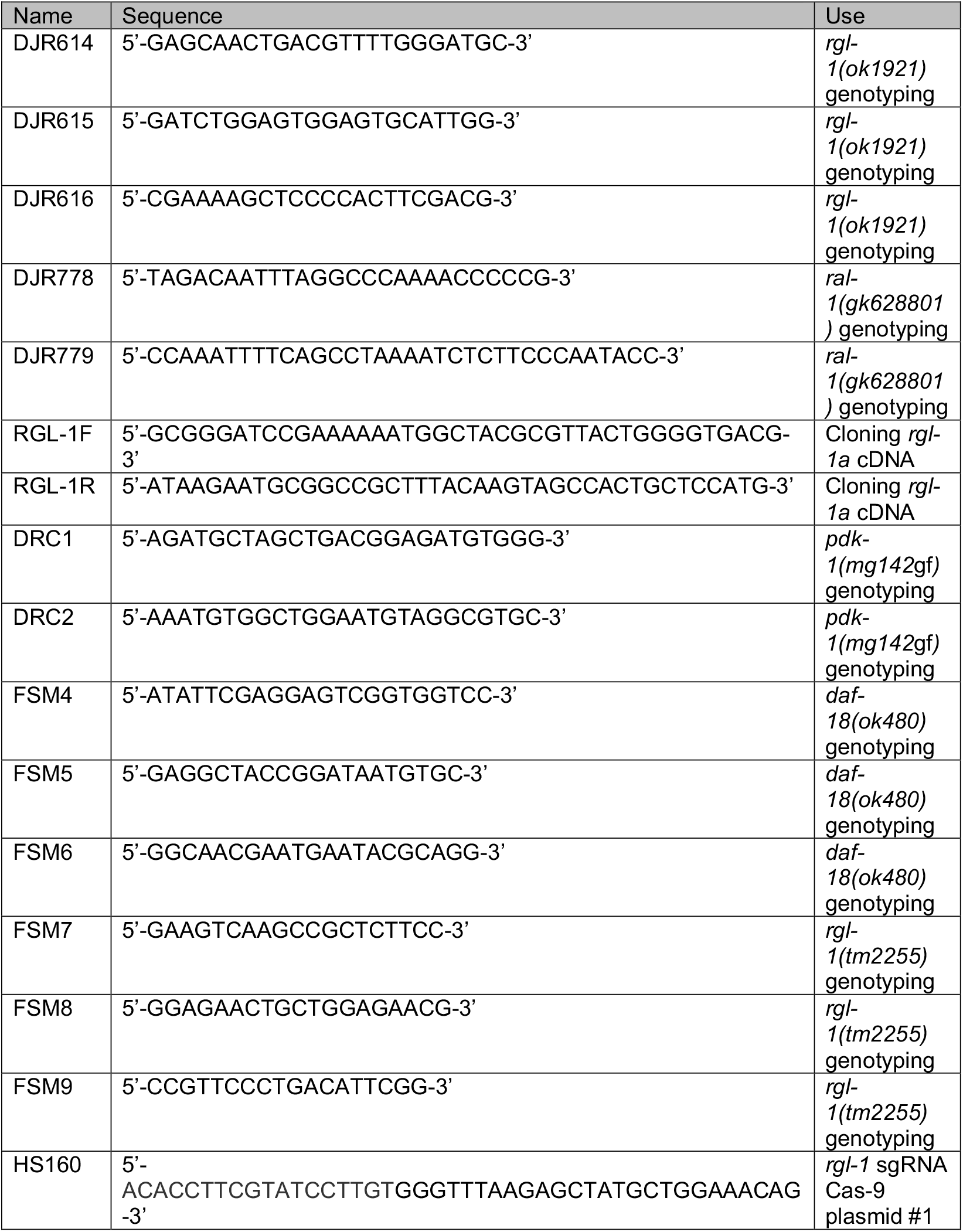
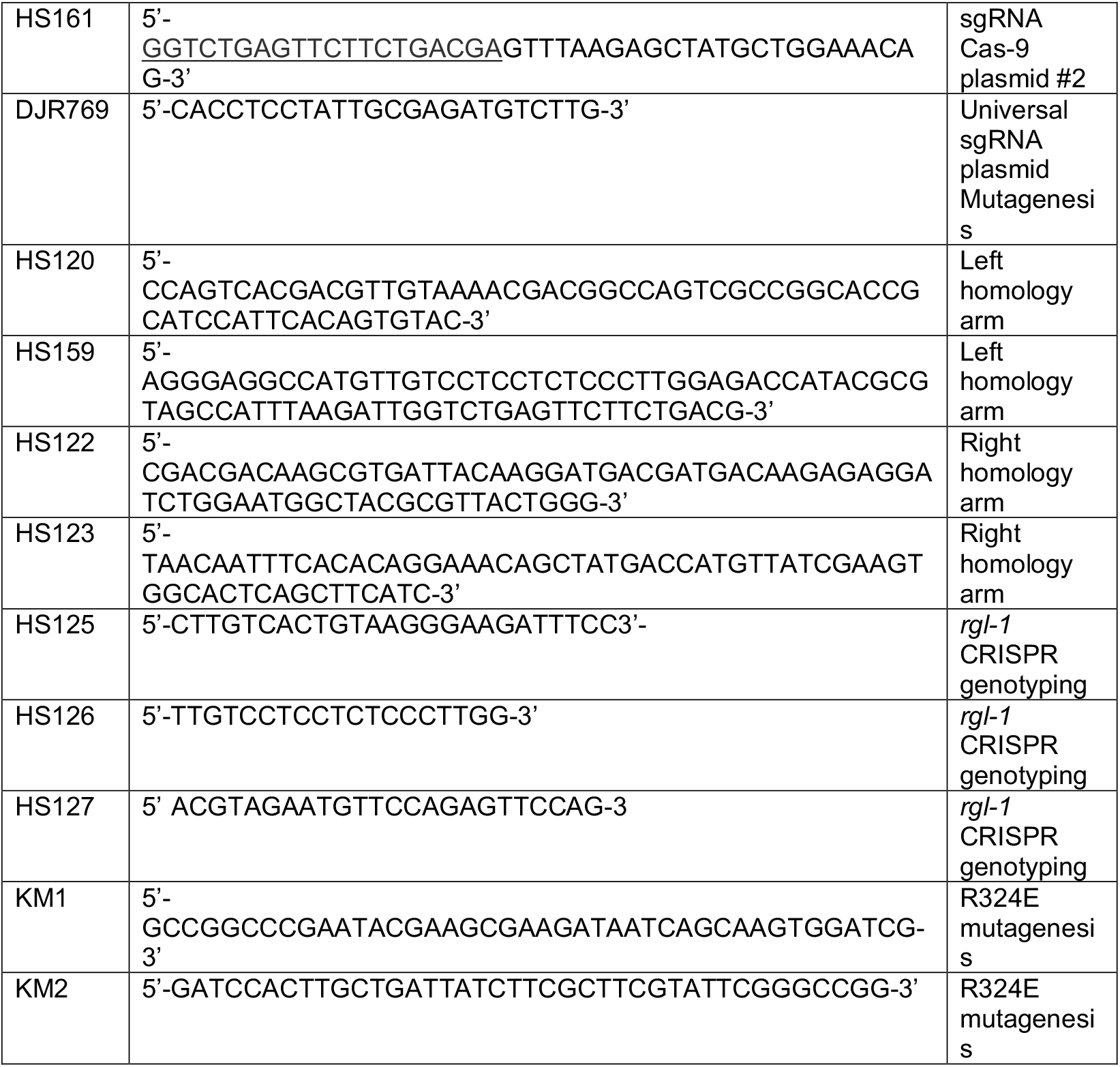
Primers

**Figure S1.**
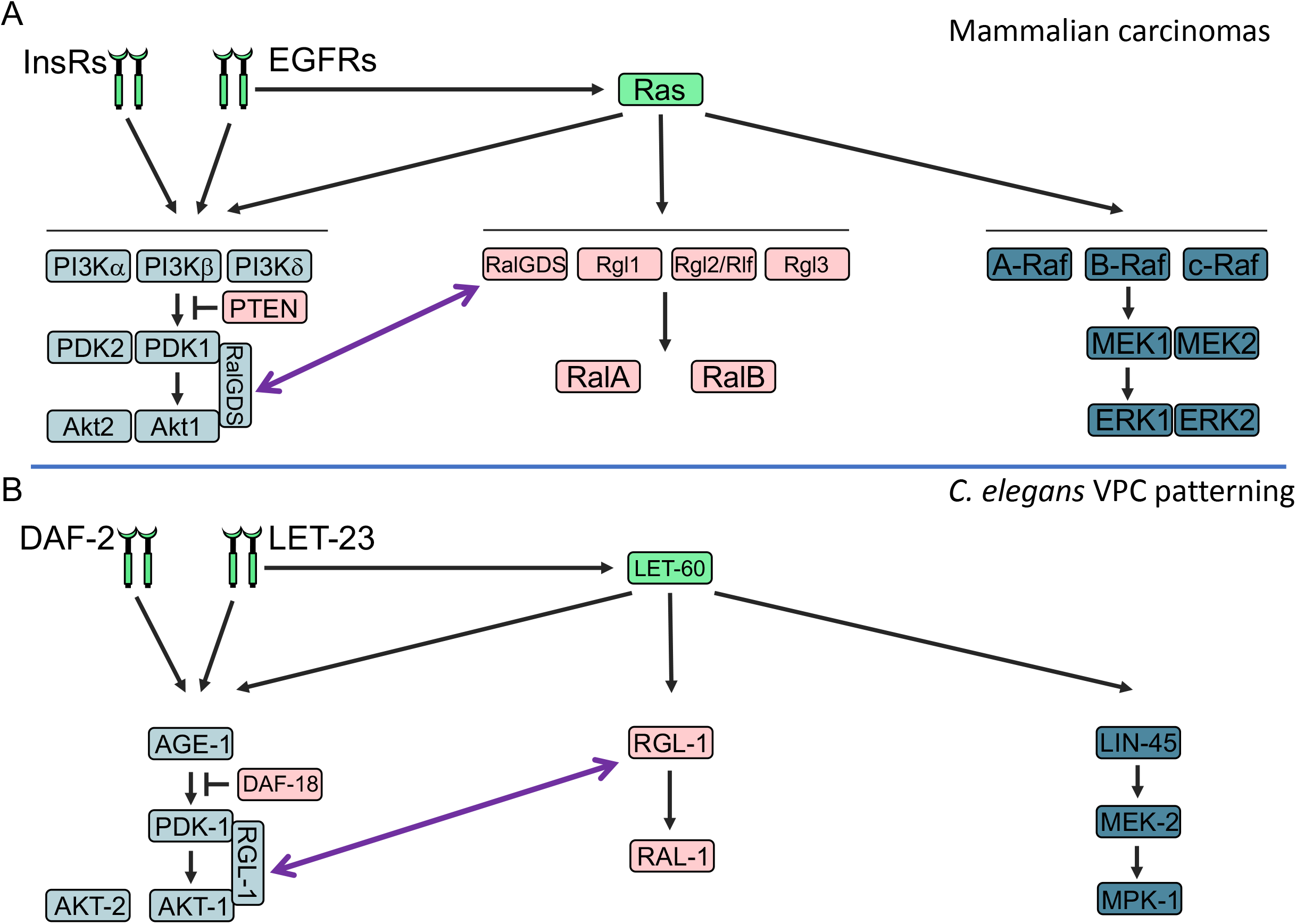
Signaling network comparisons between humans and *C. elegans.* Signaling relationships in **A)** Mammalian carcinomas and **B)** *C. elegans* VPC fate patterning. Historically, mammalian interactions have been shown directly, while *C. elegans* interactions were deduced from a combination of phenotypes, genetic epistasis, and inferences from biochemical relationships among mammalian orthologs. Color coding is the same as in other figures: blue = 1°-promoting, rose = 2°-promoter, dark = necessary and sufficient signal, light = modulatory signal. Green = both 1°- and 2°-promoting (rather than green, RGL-1 is shown in two places, with a purple two-headed arrow denoting possible dual function in both non-canonical 1°-promoting and canonical 2°-promoting roles). Activation of *C. elegans* AGE-1/PI3K by a receptor other than DAF-2, or by LET-60/Ras, has not been suggested in the literature. The interactions between RGL-1, PDK-1 and AKT-1 are inferred from genetic relationships in this study, and have not been shown directly. JIP-1, a potential intermediary between Akt and RalGDS/RalGEFs inferred from mammalian biochemical analyses in the Feig lab, is not shown.

**Figure S2.**
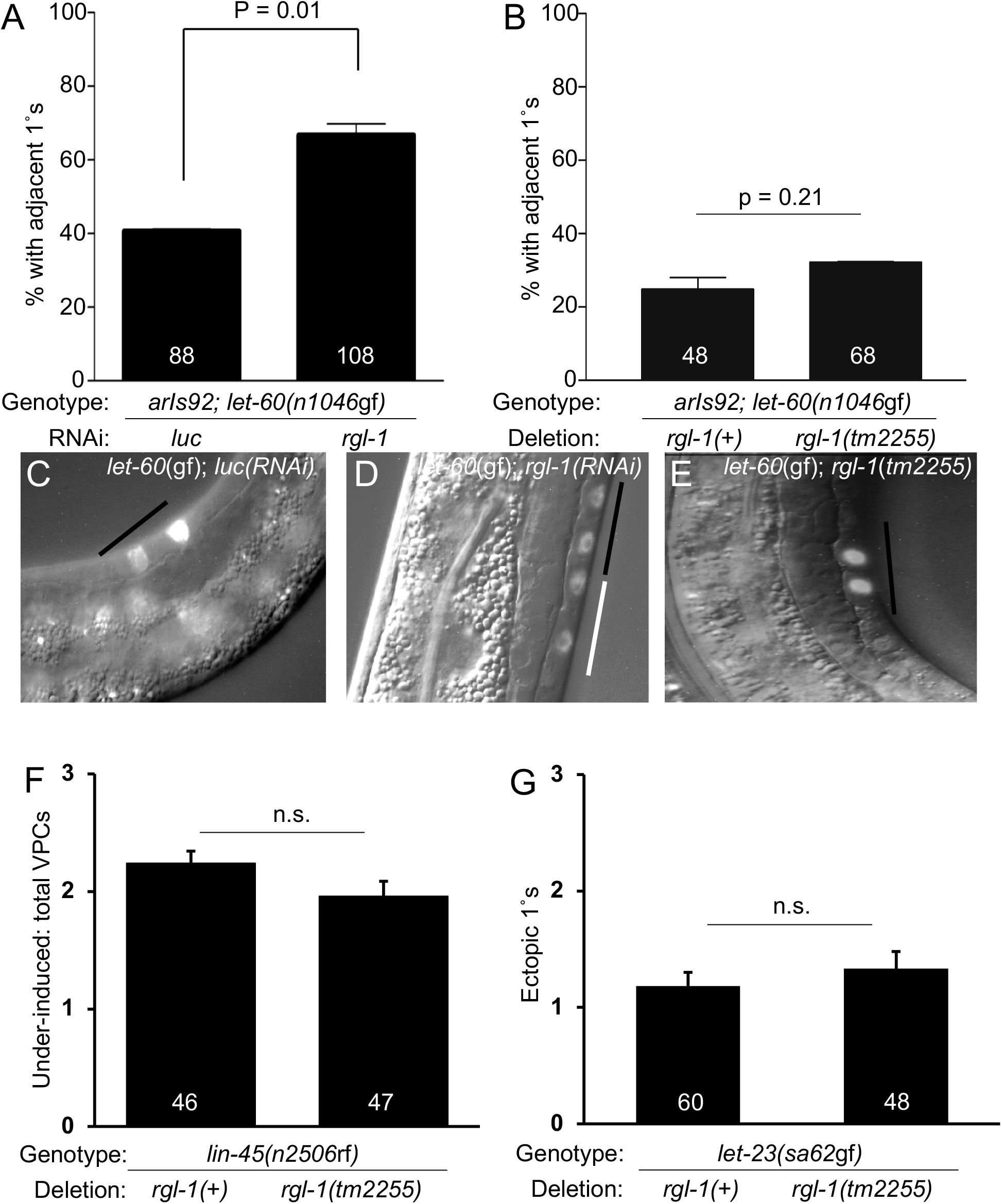
RGL-1 controls cell fate decision. **A)** Percent Pn.px-staged *let-60(gf)* L3 larvae with CFP-positive lineages neighboring the P6.p lineage (P5.p or P7.p derived) with *luc(RNAi) vs. rgl-1(RNAi)*. Shown are average percentages of animals with adjacent 1° cell fate. **B)** Percent Pn.px-staged *let-60(gf)* L3 larvae with CFP-positive lineages neighboring the P6.p lineage (P5.p or P7.p derived) with *rgl-1(+)* or *rgl-1* (tm2255Δ). Y axis is percent adjacent 1°s, white numbers in bars are number of animals scored per genotype. **C-E)** Expression of the 1 ° fate reporter *arIs92 P_egl-17_*::*cfp-lacZ* in VPC daughters. Overlaid DIC and CFP fluorescence images of **C)** *let-60(n1046gf);* luc(RNAi), **D**) *let-60(n1046gf); rgl-1(RNAi)* and **E)** *let-60(The* black bar indicates P6.px and white bar indicates P7.px cells. n1046gf); *rgl-1 (tm2255Δ)* at the Pn.px stage. **F)** Hypo-induced *lin-45(n2506)* background with and without *tm2255*. Y axis is total induced VPCs (0 = vulvaless, 3 = normal wild-type vulva.) White numbers are number of animals assayed. **G)** *let-23(sa62gf)* with and without *tm2255.* Y axis is mean ectopic 1° induction. Error bars show S.E.M. P value calculated via Mann-Whitney test.

**Figure S3.**
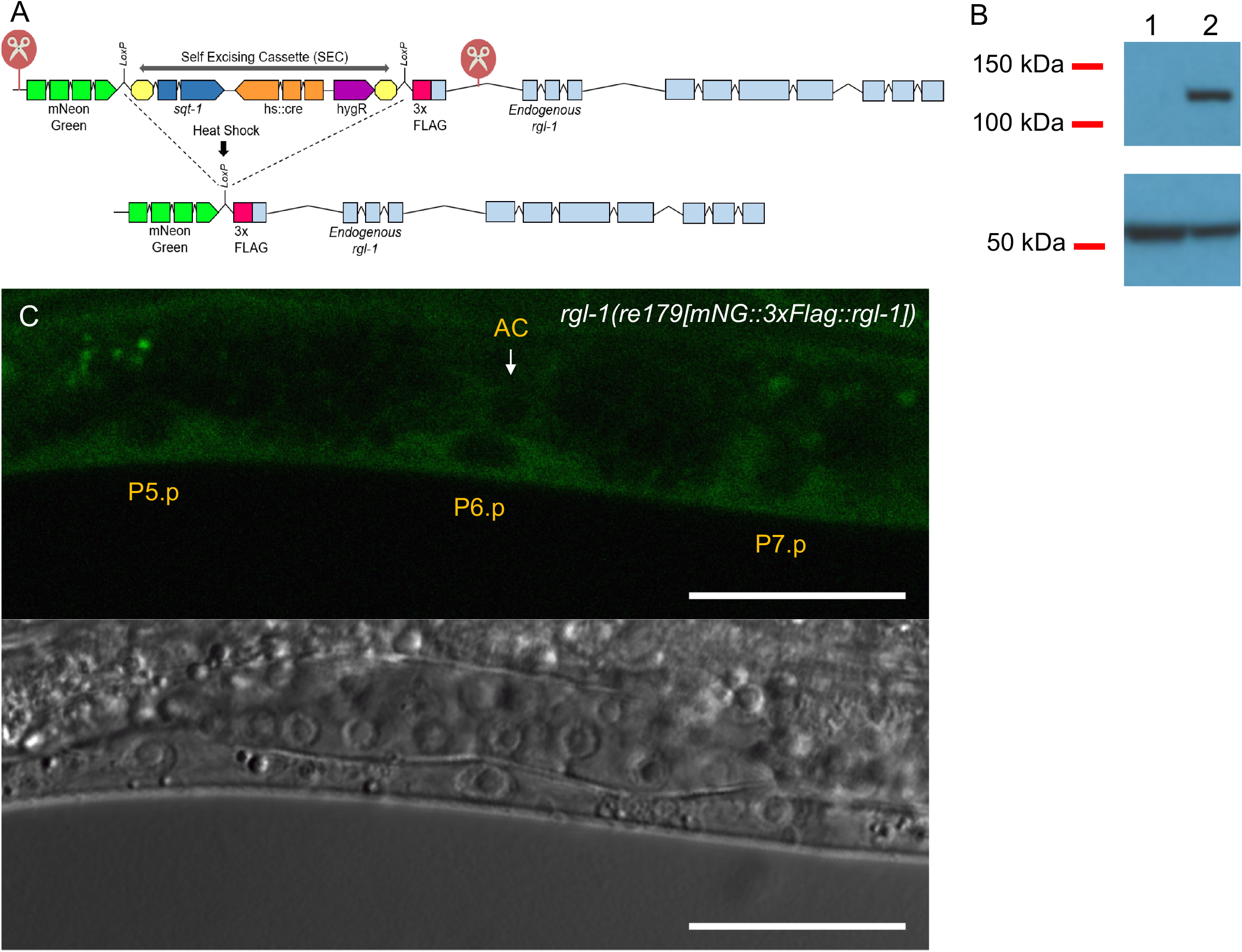
N-terminal CRISPR tagging of endogenously expressed RGL-1 protein. **A)** Strategy for N-terminal tagging of the endogenous RGL-1 protein, using the SEC approach (Dickinson, 2015). See Methods. **B)** Western blot validation of RGL-1 N-terminal tag prior to (lane 1) and after SEC excision (lane 2). Below: alpha-tubulin loading control. **C)** Tagged endogenous mNG::RGL-1 is expressed in all VPCs. Scale bar = 20 μm.

**Figure S4.**
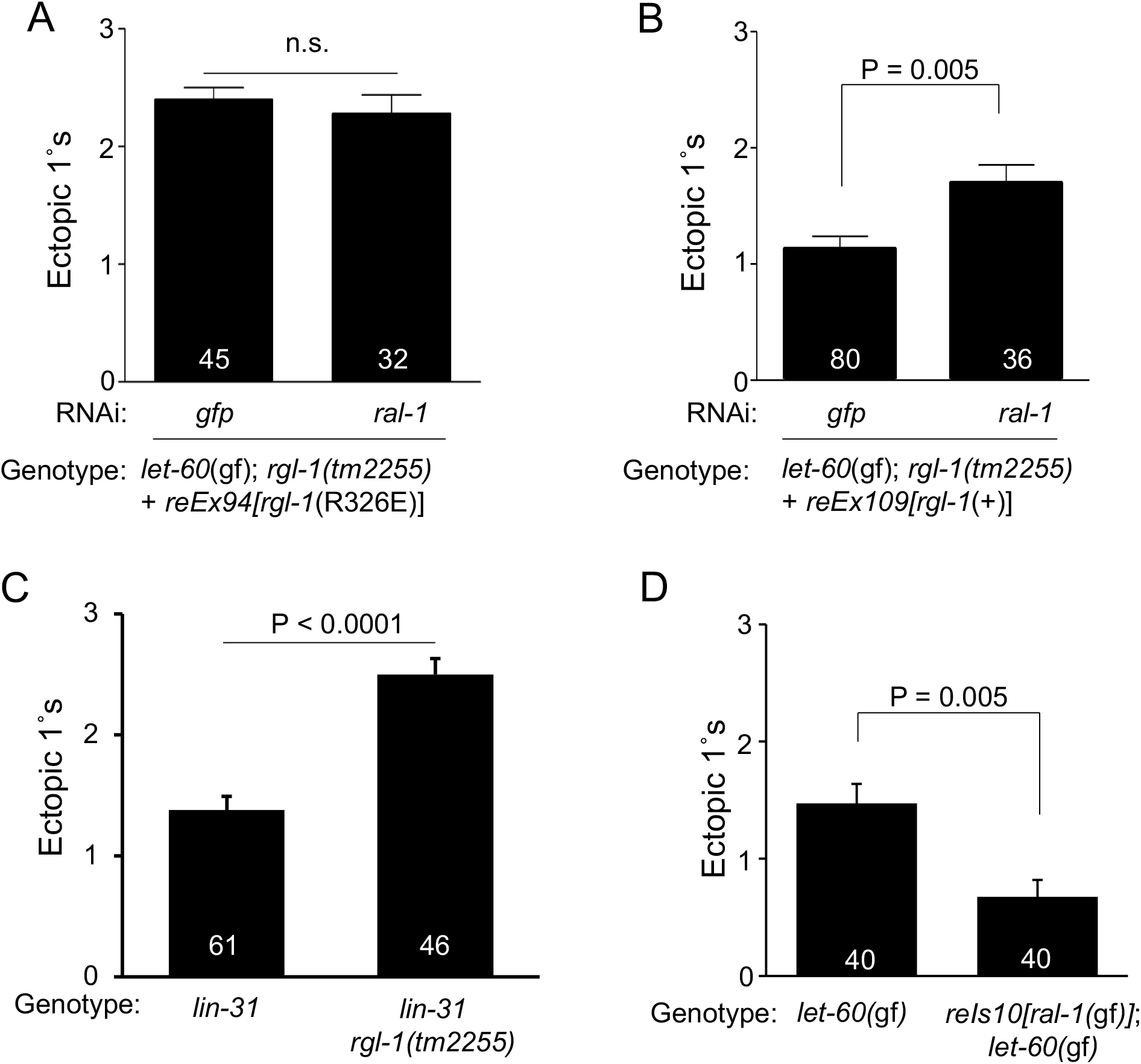
Genetically separable functions of RGL-1. **A)** *let-60{*gf); *rgl-1 (tm2255)* animals rescued by VPC-specific expression of GEF dead (R326E) RGL-1 fail to respond to *ral-1-directed vs*. control GFP RNAi. **B)** *let-* 60(gf); *rgl-1 (tm2255)* animals rescued by VPC-specific expression of wild-type RGL-1 rescue responsiveness to *ral-1*-directed but not control GFP RNAi. **C)** *lin-31(n301)* bypassed the putative Γ-promoting but not the putative 2°-promoting activity of *rgl-1,* revealed by the *tm2255* mutation enhancing ectopic 1° induction. **D)** *rels10[P_lin-31_*::*ral-1(Q75L)* + P_*myo*-2_::*gfp*] suppressed the level of ectopic 1° induction in *let-60(n1046gf,* as previously described for *reEx24* (Zand, 2011). Y axis represents mean ectopic 1° cells, white labels number of animals counted, error bars show S.E.M. P value calculated via Mann-Whitney test.

**Figure S5:**
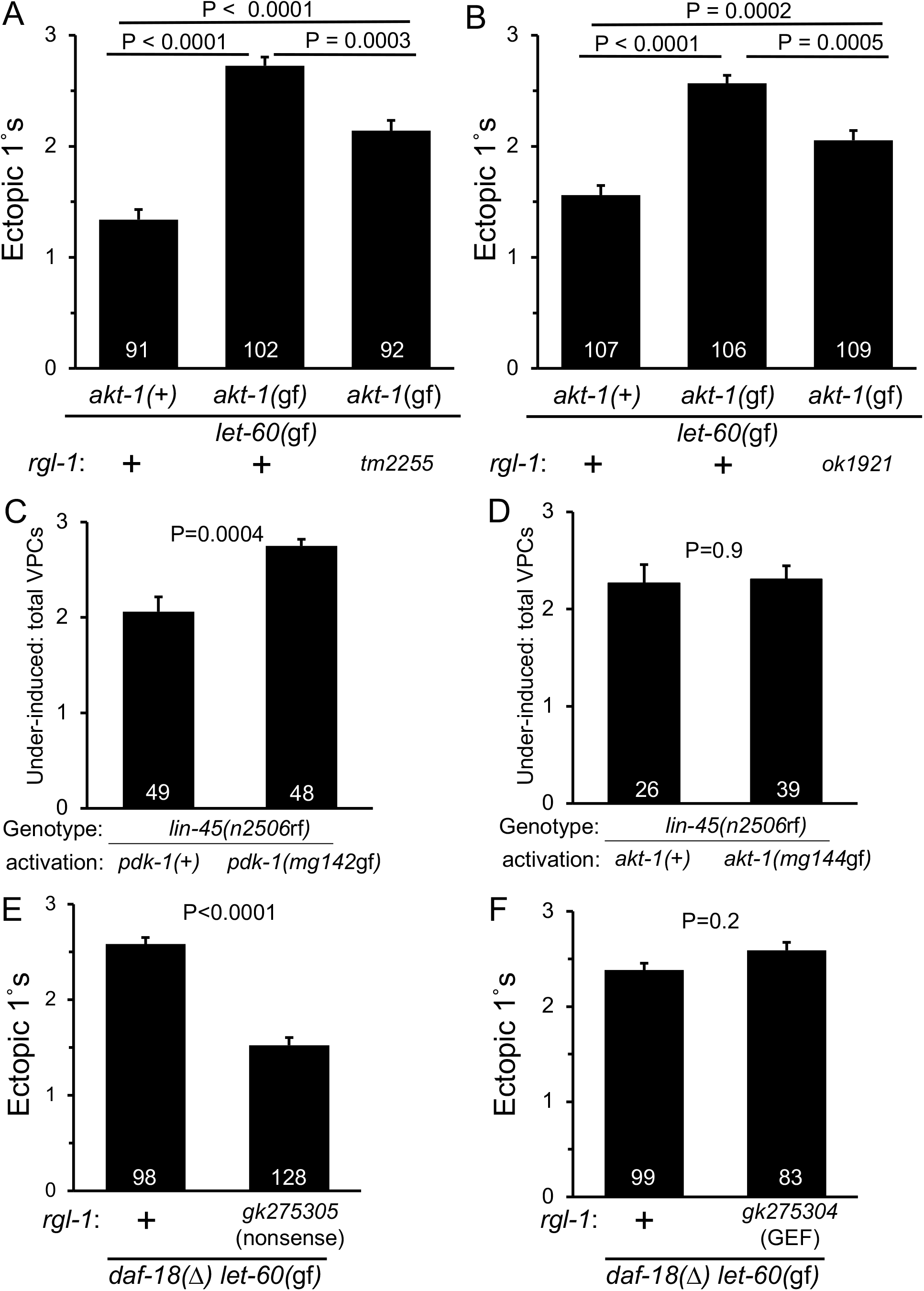
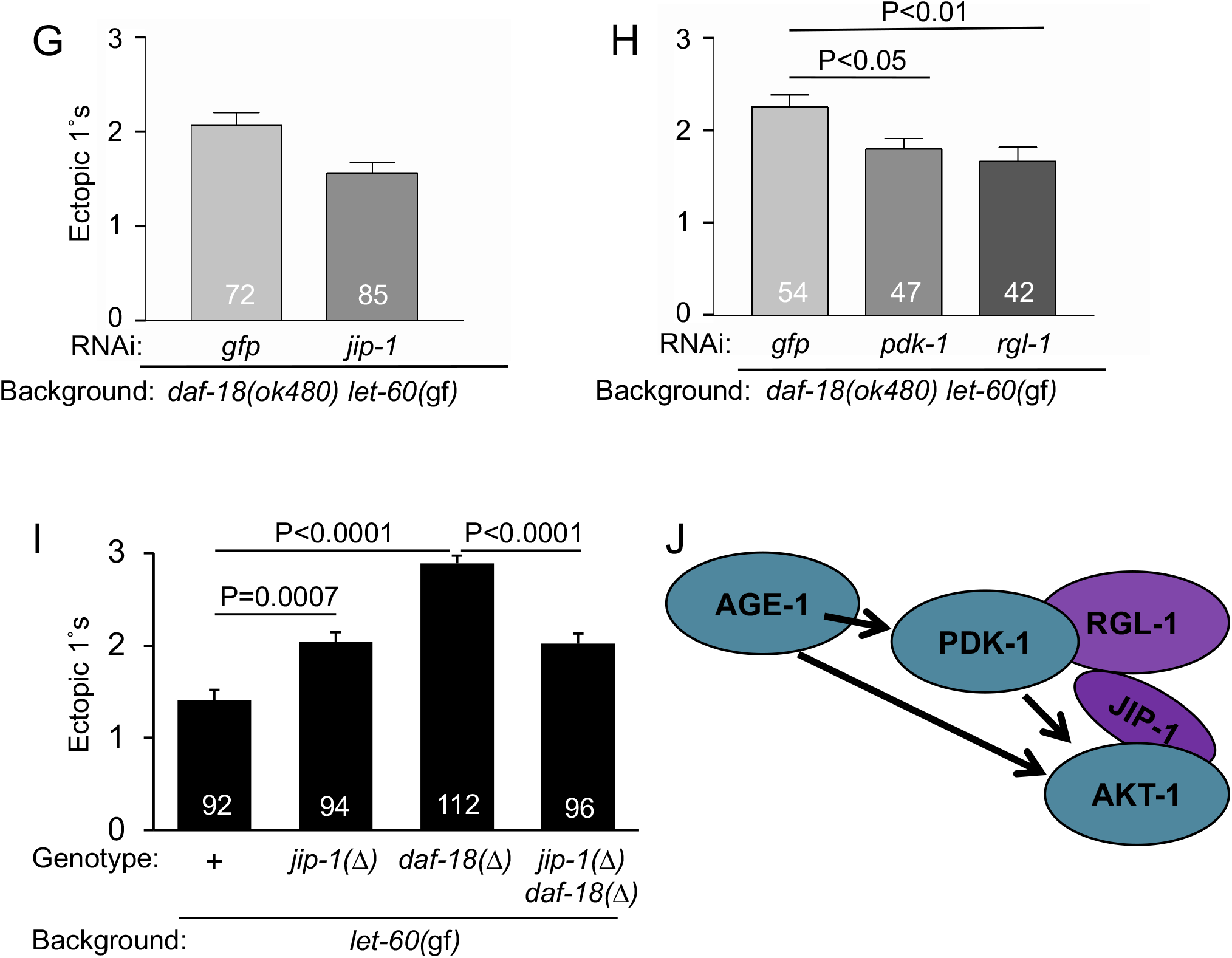
RGL-1 interacts genetically with the 1°-promoting AGE-1/PI3K-PDK-1-AKT-1 cascade. *akt-1(mg144gf)* enhancesΓ induction in *let-60(n1046gf* animals, and this enhancement is blocked by **A)** *rgí-1(tm2255)* and **B)** *rgl-1 (ok1921),* though not to the original baseline. The *lin-45(n2506)* under-induced mutant is partially suppressed by C) *pdk-1 (mg142gf* not **D)** *akt-1(mg144gf).* Unlike other assays, data shown are total induced VPCs, not ectopic Γs. *daf-18(ok480)* enhances 1° induction in *let-60(n1046gf* animals, and this enhancement is blocked by the **E)** *rgl-1 (gk275305)* nonsense mutation but not the **F)** *rgí-1 (gk275304)* R361Q putative GEF dead mutation. **G)** y/p-7-directed RNAi suppressed the increase in ectopic 1° induction in the *íet-60(n1046gĩ)* background conferred by *daf-18(ok480),* compared to *gfp(RNAi)*. **H)** *pdk-1-* and *rgl-*7-directed RNAi similarly suppress *daf-18(ok480) let-60(n1046gf* ectopic 1° induction. I) The *jip-1* deletion mutation, *jip-1(tm6137),* enhances *let-60(n1046gf* alone but suppresses *daf-18(ok480) let-60(n1046gf,* consistent with JIP-1 performing two functions. **J)** The observed genetic interactions are consistent with RGL-1 and JIP-1 functioning in the AGE-1-PDK-1-AKT-1 1° promoting cascade, as described for mammalian orthologs (Tian *et al.,* 2002; Hao Wong and Feig, 2008). Except for *lin-45(rf),* Y axis represents mean ectopic 1° cells, white labels number of animals counted, error bars show S.E.M. P value calculated via Mann-Whitney test or ANOVA.

**Figure S6:**
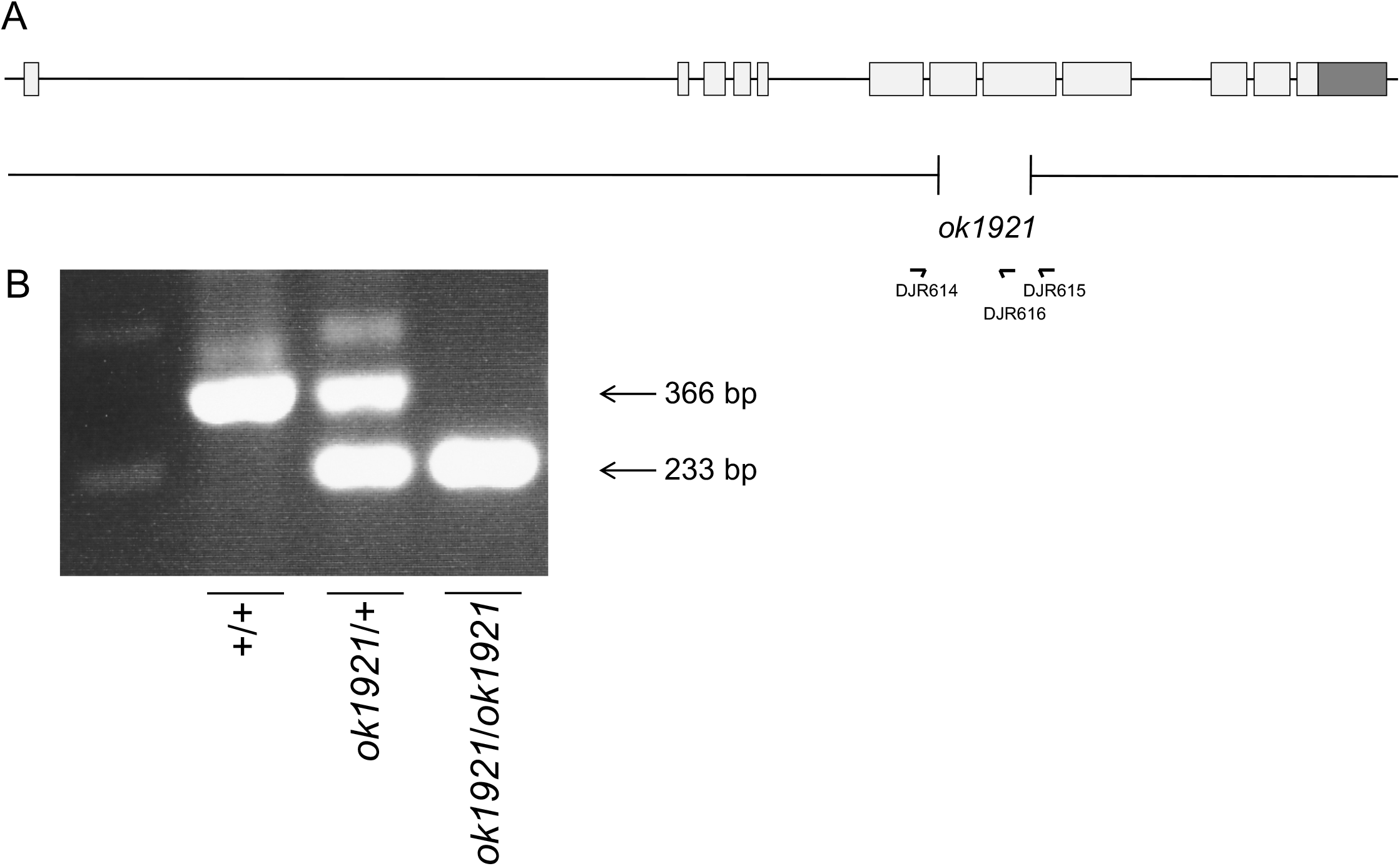
PCR detection of *rgl-1(ok1921).* **A)** A scale schematic of the *rgl-1* gene, the *ok1921* lesion, and detection primers. **B)** Agarose gel of +/+, *ok1921l+* and *ok1921/ok1921* single animal PCR reactions.

**Figure S7:**
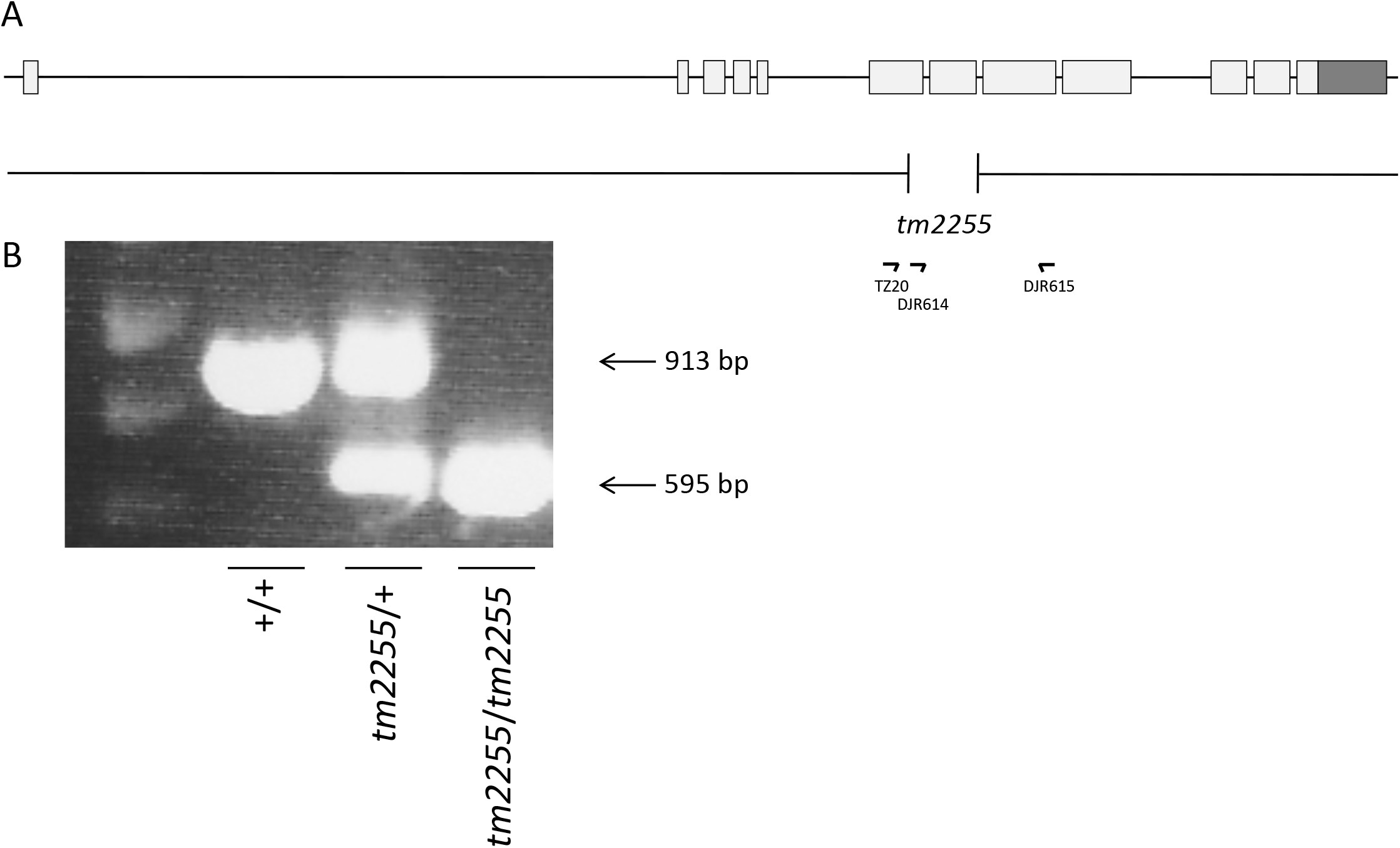
PCR detection of *rgl-1(tm2255).* **A**) A scale schematic of the *rgl-1* gene, the *tm2255* lesion, and detection primers. **B**) Agarose gel of *+/+, tm2255/+* and *tm2255/tm2255* single animal PCR reactions.

